# Identifying tissue states by spatial protein patterns related to chemotherapy response in triple-negative breast cancer

**DOI:** 10.1101/2025.10.06.680783

**Authors:** L.M. Luque, A. Khan, G. Torrisi, Tessa D. Green, D. Hardman, C. Owczarek, T. A. Phillips, D.S. Marks, M. Parsons, C. Sander, L.J. Schumacher

## Abstract

Triple-negative breast cancer (TNBC) is an aggressive malignancy with limited targeted therapies and variable responses to conventional chemotherapy, influenced by intratumoral heterogeneity and complex tumor microenvironment (TME) interactions. Understanding spatiotemporal cellular interplay and tissue organization is crucial for advancing tumor biology and improving patient stratification. Spatially resolved proteomics, such as Imaging Mass Cytometry (IMC), offers a powerful approach to dissect the TME. We present an end-to-end computational pipeline for robust quantitative analysis of large-scale IMC datasets, addressing the challenge of batch effects through image-level contrast adjustment. Applying this framework to 813 tissue regions encompassing over 4 million cells from 63 TNBC patients, we revealed distinct spatial arrangements of cell types between chemotherapy responders and non-responders. Non-responders showed reduced cytotoxic T-cell infiltration into tumor regions and increased spatial co-localization between fibroblasts and macrophages, a pattern that persisted and intensified after chemotherapy treatment. To integrate these complex spatial-molecular relationships, we used graph neural networks (GNNs) to predict treatment response from pre-treatment samples with AUROC=0.71. Interpretability analysis identified B7H4, CD11b, CD366, and FOXP3 as the most predictive protein markers, with fibroblasts, cancer cells, and CD8+ T cells being the most informative cell types. This study introduces a scalable analytical framework for spatial proteomics with interpretable predictions, suggesting features of tissue state that could guide treatment decisions in TNBC and further our understanding of the spatial determinants of therapeutic response.

## Introduction

### Challenges in TNBC treatment

Breast cancer is a globally prevalent disease, posing considerable challenges in treatment and management due to high incidence and mortality rates worldwide (Sung et al. 2021). Among its subtypes, triple negative breast cancer (TNBC) is particularly aggressive, characterized by the lack of estrogen receptor (ER), progesterone receptor (PR), and human epidermal growth factor receptor 2 (HER2) (Bianchini et al. 2016; Mukund et al. 2025). ER, PR and HER2 are molecular targets for specific targeted therapies in breast cancer; their absence in TNBC means that these targeted therapies, such as anti-estrogen (e.g., tamoxifen) or anti-HER2 (e.g., trastuzumab) drugs, are ineffective for treating TNBC patients (Shetti et al. 2019). With limited targeted therapeutic options, TNBC treatment has largely relied on conventional chemotherapy, such as anthracyclines and taxanes, most commonly as neoadjuvant chemotherapy (NACT). However, there is significant variability in the response rates to chemotherapy, resulting in unnecessary toxicity for non-responders (Bianchini et al. 2016; Mukund et al. 2025; X. Wang et al. 2024). Predicting a patient’s response to chemotherapy is crucial for personalized treatment planning, enabling the identification of alternative treatments and minimizing harm due to ineffective treatment.

### IMC in TNBC profiling

Recent advancements in spatially resolved imaging technologies, such as Imaging Mass Cytometry (IMC), have made it possible to quantify multiple protein markers at the resolution of individual cells, enabling detailed interrogation of the tumor microenvironment. IMC uses metal-tagged antibodies to detect and quantify up to 40 proteins or other molecules in biological samples, with a spatial resolution of 1 µ𝑚 and an ablation frequency of 200 Hz (Giesen et al. 2014). This technology offers a powerful tool to spatially profile regions of interest (ROIs) in patient biopsies, enabling analysis of tumour microenvironments and characterization of pathological features in diseases such as TNBC. These capabilities open new avenues for designing precision cancer therapeutics by revealing spatial correlates of immune evasion, therapy resistance, and disease progression. Most prior studies have focused on relatively small datasets (tens of ROIs), which often allows for manual, semi-supervised workflows but limits generalizability (Milosevic 2023; Windhager et al. 2023). However, analysis of IMC data at scale presents challenges: Larger datasets introduce heterogeneity in batch effect and imaging conditions over time which can complicate data preprocessing and cell type annotation.

### End-to-End IMC analysis pipeline

Analysis of IMC data involves many of the same difficulties inherent to other single-cell modalities, specifically the difficulty of distinguishing noise and sample-to-sample or ROI-to-ROI variation. The field has not yet settled on a single best practices workflow, and most analysis relies on a mix of ad hoc procedures and more finalized packages (reviewed in (Milosevic 2023)). Here we present an end-to-end pipeline that integrates existing analysis tools with new approaches to overcome specific challenges in the analysis of large and heterogeneous IMC datasets. The pipeline covers pre-processing of raw IMC data including normalisation and batch-correction, annotation of cell types, their abundance, and spatial organisation, and prediction of response to therapy.

### TNBC IMC dataset

To demonstrate the applicability of this workflow to large-scale datasets, we investigate spatial determinants of chemotherapy response in TNBC by analyzing a recently created Imaging Mass Cytometry (IMC) dataset (Parsons et al. 2025). This IMC dataset was generated as part of the SMART multimodal atlas <u>(Wall et al. 2025)</u>, which combines spatial transcriptomics, histological analysis, and IMC data from the same patient cohort. The IMC dataset includes over 4 million cells derived from 813 regions of interest (ROIs) across 63 patients, incorporating both pre-treatment biopsies and post-treatment resections from non-responders. All patients received EC-T or EC-T-carboplatin neoadjuvant chemotherapy regimens. The tissue samples were stained with a marker panel targeting 35 tumor, immune, and stromal proteins. An overview of the dataset and the analytical workflow is provided in Figure 1, which illustrates the study design, number of ROIs per response class, total cell count, chemotherapy regimen, and the end-to-end image analysis pipeline.

**Figure 1.**
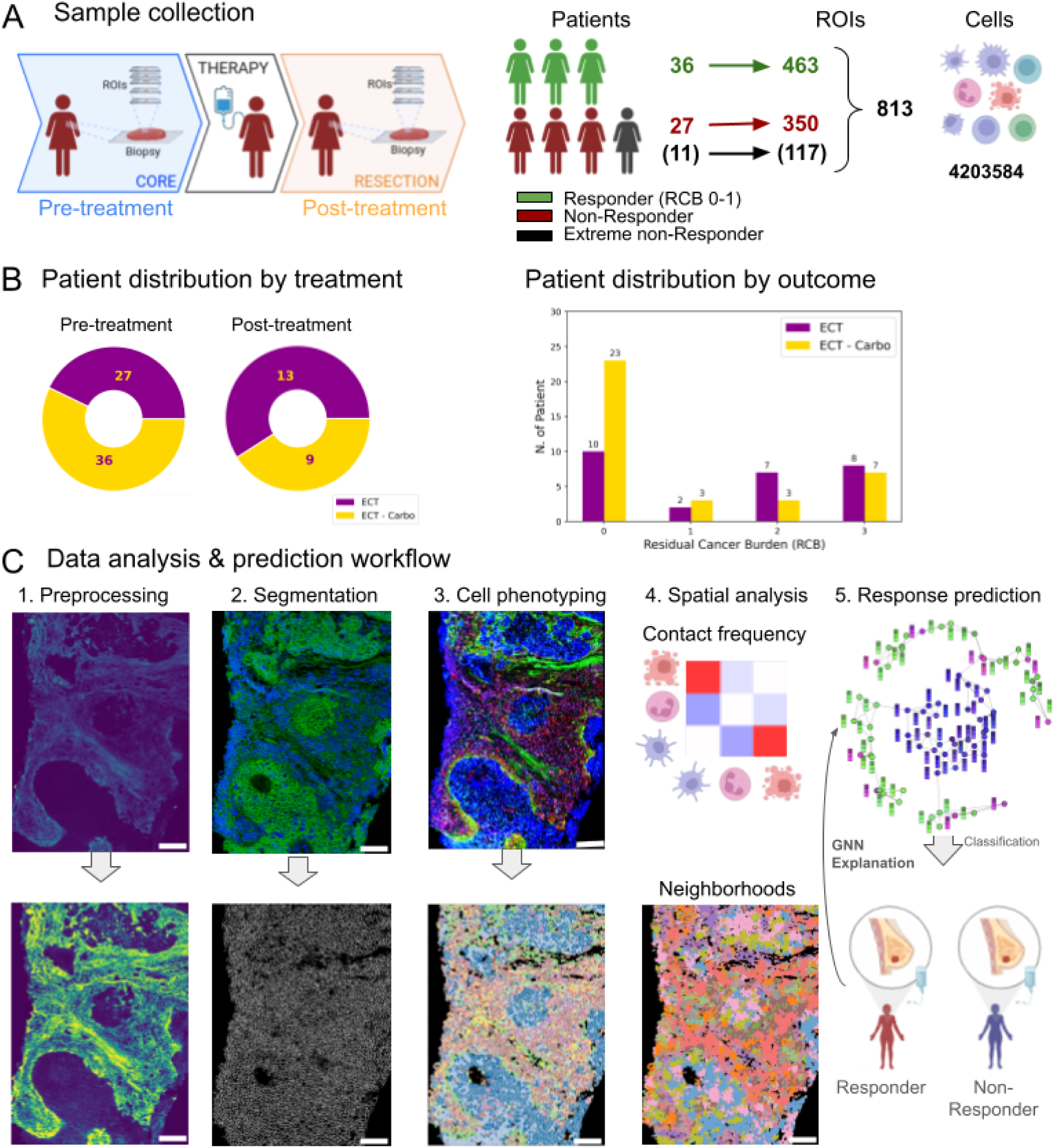
Overview of dataset and analysis workflow. **A** Biopsies were taken before and after NACT (left), and multiple regions of interest (ROIs) extracted from each biopsy, stained with metal-tagged antibodies and imaged using Imaging Mass Cytometry (IMC). Number of ROIs in each response class (middle) and total number of cells identified in the dataset (right). **B** All patients received EC-T chemotherapy, and some patients with carboplatin (EC-T-carbo (left). Distribution of patients over residual cancer burden (right). **C.** Analysis workflow for IMC data showing an illustrative example of preprocessing, segmentation, cell phenotype, spatial analysis, and response prediction. The scale bar is 150µm. Image colours show the following: 1. Preprocessing: intensity for representative IMC channel. 2. Segmentation: Representative IMC channels in green and blue (top) and segmentation mask (bottom). 3. Cell phenotyping: Various IMC channels used for cell phenotyping (top) and cell type labels after segmentation (bottom). 4. Spatial analysis: schematic heatmap illustrating high (red) or low (blue) co-localisation between pairs of cell types (represented by cartoon drawings) (top) and cells coloured by neighborhood cluster label (bottom). 5. Response prediction: cells are nodes on the graph, edges represent contact between nearest neighbors, and each cell also carries a feature vector (here represented by columns) of protein expression and/or cell type label.

Complementary to IMC, spatial transcriptomics approaches such as those used in the parallel SMART study (Wall et al. 2025) can capture the molecular signatures underlying spatial cell type patterns, providing orthogonal validation and complementary mechanistic insights into the tumor-immune environments, while our analysis emphasises the spatial organisation and cell-cell interactions revealed through single-cell resolution protein expression.

### Image-level batch correction

As IMC and other multiplexed protein imaging technologies become more widely used and dataset sizes increase, batch correction becomes more and more important. One commonly used end-to-end workflow (Windhager et al. 2023) relies on batch correction methodologies originally developed for RNA-seq data, which means that any errors in cell segmentation and earlier analysis bleed over into the batch correction procedure. Here, we use contrast-limited adaptive histogram equalisation (CLAHE) (Pizer et al. 1987) for batch correction on the level of individual images prior to cell segmentation and subsequent analyses.

### Prediction of chemotherapy response

Previous analysis of IMC data from another TNBC cohort has shown selected spatial features, including cell density, proliferative fractions, and cell-cell contacts can be used to predict response to immunotherapy but not chemotherapy alone (X. Q. Wang et al. 2023). To test the predictive information in pre-treatment samples beyond selected spatial features, we use the cell-contact graph with all protein channels as input for machine learning to predict chemotherapy response. Graph neural networks (GNNs) have demonstrated promising capabilities in capturing spatial relationships and modeling complex biological networks (Xu* et al. 2018; Li, Huang, and Zitnik 2022). GNNs are inherently designed to handle graph-structured data, making them a suitable candidate to integrate protein abundance and celltype labels (nodes), and cell-cell contact information (edges) to learn features that capture disease state and progression. We train a GNN model on pre-treatment samples to predict response to chemotherapy and to interpret which signals are most important for the prediction.

## Results

### Contrast adjustment to mitigate against technical variability

We observed systematic differences in average protein count, likely due to batch effects that occurred through samples being stained and imaged in groups, and variation in sample quality (Fig. S2A). The batch correction problem is well known in single cell transcriptomics, and several tools have been developed (Chu et al. 2022). Some of these tools from the single cell transcriptomics field have been used in IMC analysis pipelines (Windhager et al. 2023) to correct for batch effects. However, many scRNAseq batch correction methods are specifically developed for the statistics of sequencing-based mRNA counts, not image-based protein measurements. Furthermore, methods that apply batch correction at the cell level rely on cell segmentation being processed first, but cell segmentation could in turn be sensitive to batch effects. We instead opted for an approach that corrects for batch correction at the image level, using contrast-limited adaptive histogram equalisation (CLAHE; Fig. 2A, see Methods). This successfully removed batch effects, at least in lower dimensional visualisations (Fig. 2B & S2A). After denoising and batch-correction, we proceeded with cell segmentation (Fig. 2C, see Methods).

**Figure 2.**
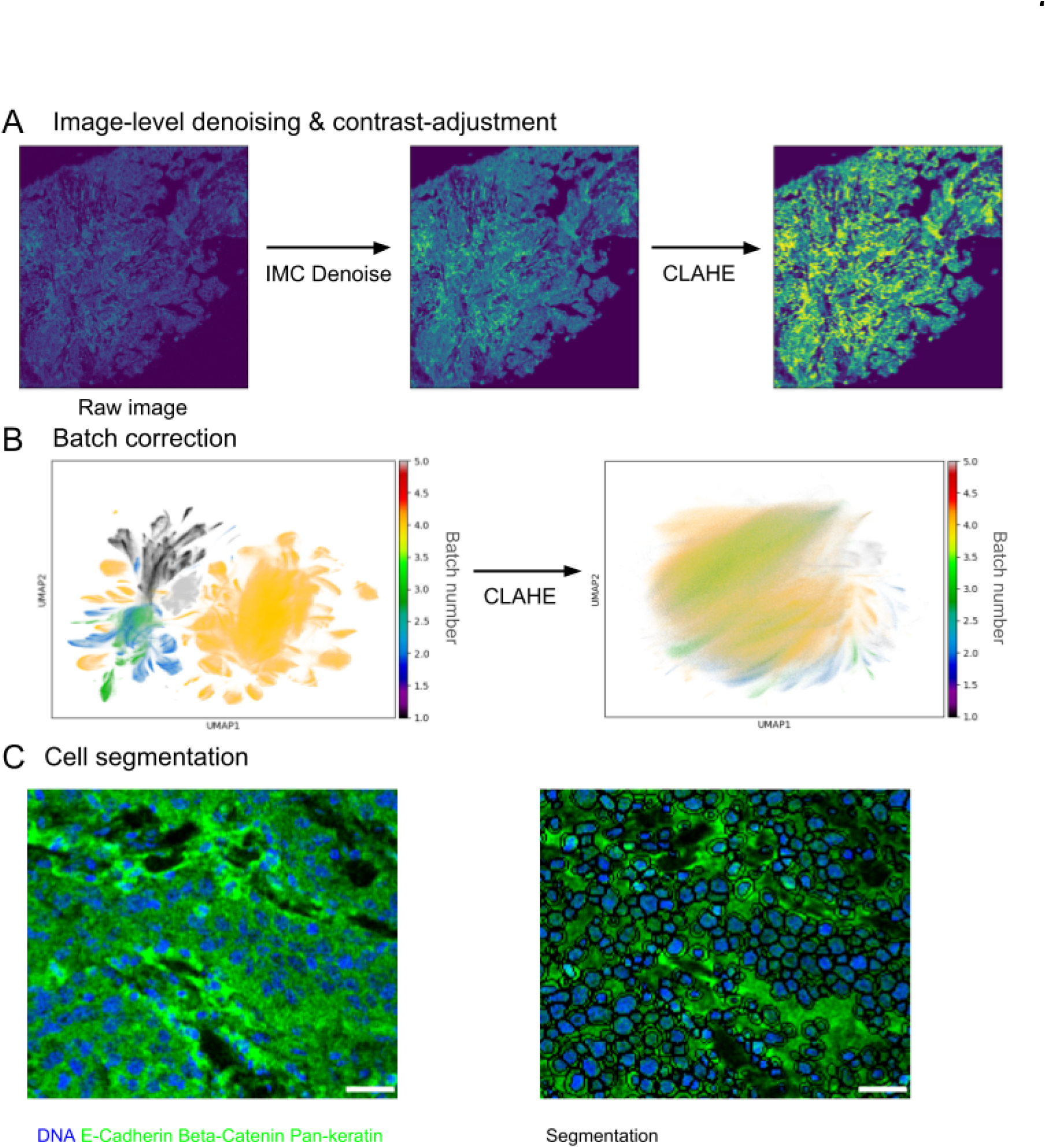
Data preparation: denoising, batch effects, segmentation. **A** Denoised and contrast enhanced image of a representative ROI. **B** Approximate overlap of data from different patients reflects removal of batch effects. Each point is a single cell. After histogram equalisation (CLAHE) the batches are no longer separated. **C** Representative raw image (left) and with overlaid segmentation boundaries for nuclei and whole cells (right). Blue: DNA antibody; Green: Combination of E-Cadherin, Beta-Catenin, and Pan-keratin staining.

### Cell type annotation resolves 18 cell clusters

We annotated cells with Pixie (C. C. Liu et al. 2023), which uses unsupervised clustering of pixels to annotate cell-level features for clustering and subsequent annotation of cell types and states (see Methods). Of the 35 markers in the original panel (Fig. S2B), we used a subset for annotating cell types (Fig. 3A), with Ki-67 as an indicator of proliferation status (see Methods). This approach identified 18 cell clusters (Fig. 3A). To differentiate further between proliferative and non-proliferative cancer cells, we split both cancer clusters based on their Ki67 abundance (Fig. 3B, Fig. S3-1B). We mapped the resulting clusters back onto the images (Fig. 3C) and analysed the abundance and spatial arrangement of cell types across samples. The annotated cell types show the heterogeneous spatial structure of biopsies, typically with regions rich in cancer cells, and regions rich in immune cells (Fig. 3C). Overall, the most abundant cell types were CD8+ T cells, cancer cells, and fibroblasts (Fig. 3D), with approximately one third of all cancer cells marked as proliferative based on Ki67 expression (Fig. 3E).

**Figure 3.**
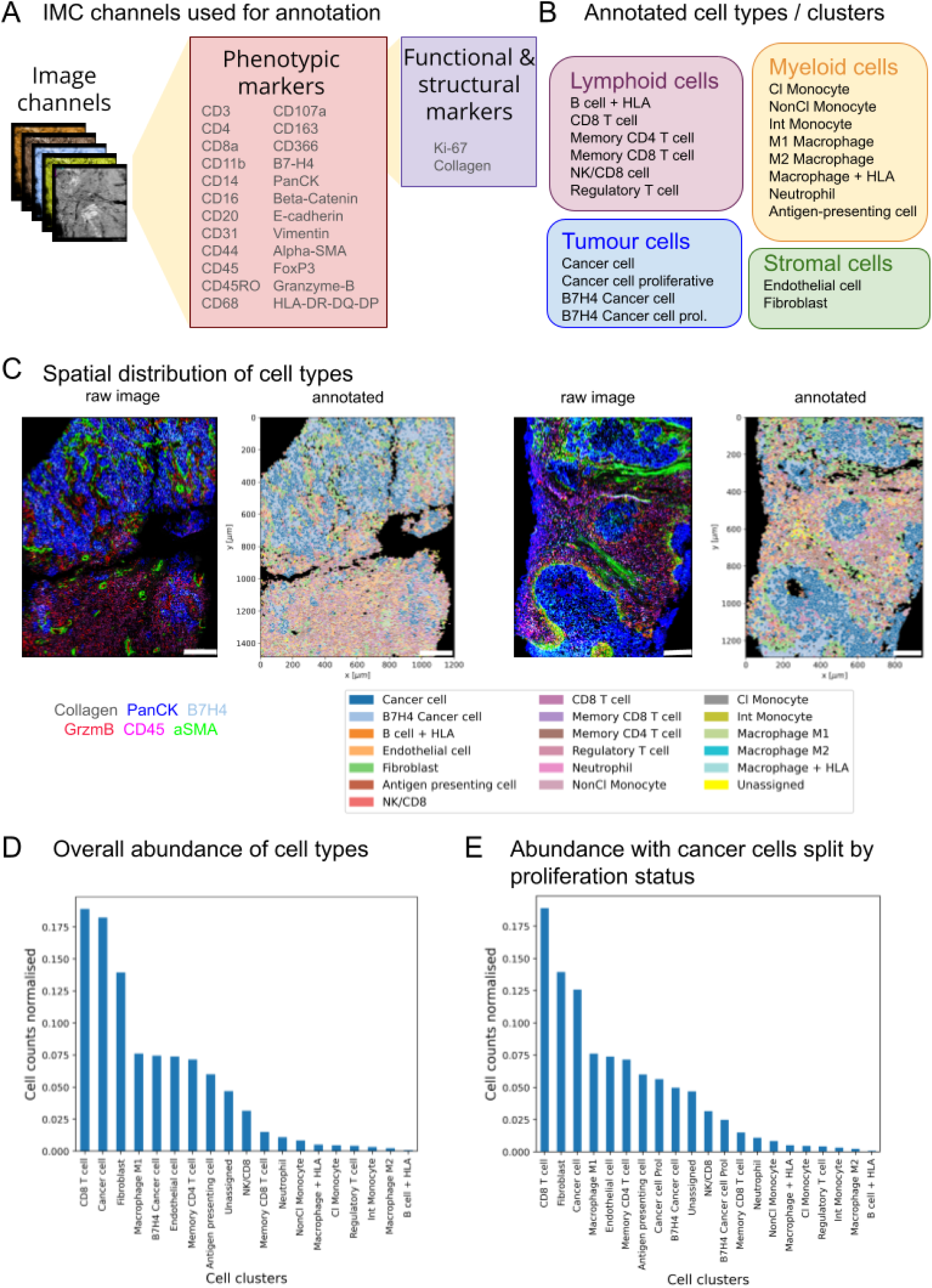
Cell type assignment and abundance. **A** IMC channels used for annotation of cell type (phenotypic markers) and functional state (functional markers). **B** Cell clusters annotated. **C** Representative raw images of selected channels (Collagen, PanCK, B7H4, GrzmB, CD45, αSMA) and corresponding cell type labels (see legend). The scale bars are 200µm on the left and 150µm on the right. **D** Relative abundance of cell clusters across all ROIs, without subdividing proliferating cells. **E** Relative abundance of cell clusters across all ROIs, with sub-divided Cancer cell and B7H4 Cancer cell clusters into proliferation and non-proliferating based on Ki67 expression.

### Spatial organization of cell types in responder and non-responder tissues

We investigated tumor-immune microenvironments, and relationship to patient chemotherapy response. Antigen-presenting cells, Non-Classical Monocytes, and Regulatory T cells were more abundant in responders, and CD8+ T cells and M2 macrophages more abundant in non-responders (Fig. S4-1A,B), all with small effect sizes. Cancer cells expressing B7H4 are reported to be immunosuppressive (Zhou et al. 2023) and associated with worse prognosis in TNBC (L. Wang et al. 2018). The proportion of B7H4+ cancer cells as a function of total cancer cells, irrespective of proliferation status, was not statistically significant between responder and non-responer patient tumours (Fig. S4-1C). However, the respective fraction of Ki67+ B7H4+ and B7H4-cancer cells were higher in responders (Fig. S4-1D), consistent with increased proliferative index correlating with chemotherapy sensitivity.

Neighbourhood enrichment analysis revealed several pairs of cell types to be in contact more or less frequently than in a random re-arrangement of cells, and some pairs of cell types less frequently (Fig. S4-2A&B). CD8+ T cells co-locate more with CD4+ memory T cells in responders (Fig. 4A), consistent with CD4+ memory T cells supporting CD8+ cytotoxic T cell activity to kill the tumour. B7H4 expressing cancer cells were on average not more often in contact with other cancer cells in responders than non-responders, with only a subset of ROIs in responders with higher enrichment constituting a longer tail in the distribution of neighborhood enrichment z-scores (Fig. 4A). In contrast, cancer cells were less often in contact with macrophage+HLA cells in responders (Fig. 4A). This could indicate that antigen-presenting (HLA-DR+) macrophages are spatially excluded from tumour regions in responders, limiting the direct tumour-macrophage crosstalk that can drive immune evasion (Tharp et al. 2024). CD8+ T cells and proliferative cancer cells also co-located more in non-responders (Fig. 4A). Although counter-intuitive, it is possible that proliferating tumour regions secrete chemokines which recruit CD8 T cells, but those cells could be unable to exert cytotoxic effects due to T cell exhaustion or metabolic restriction.

**Figure 4.**
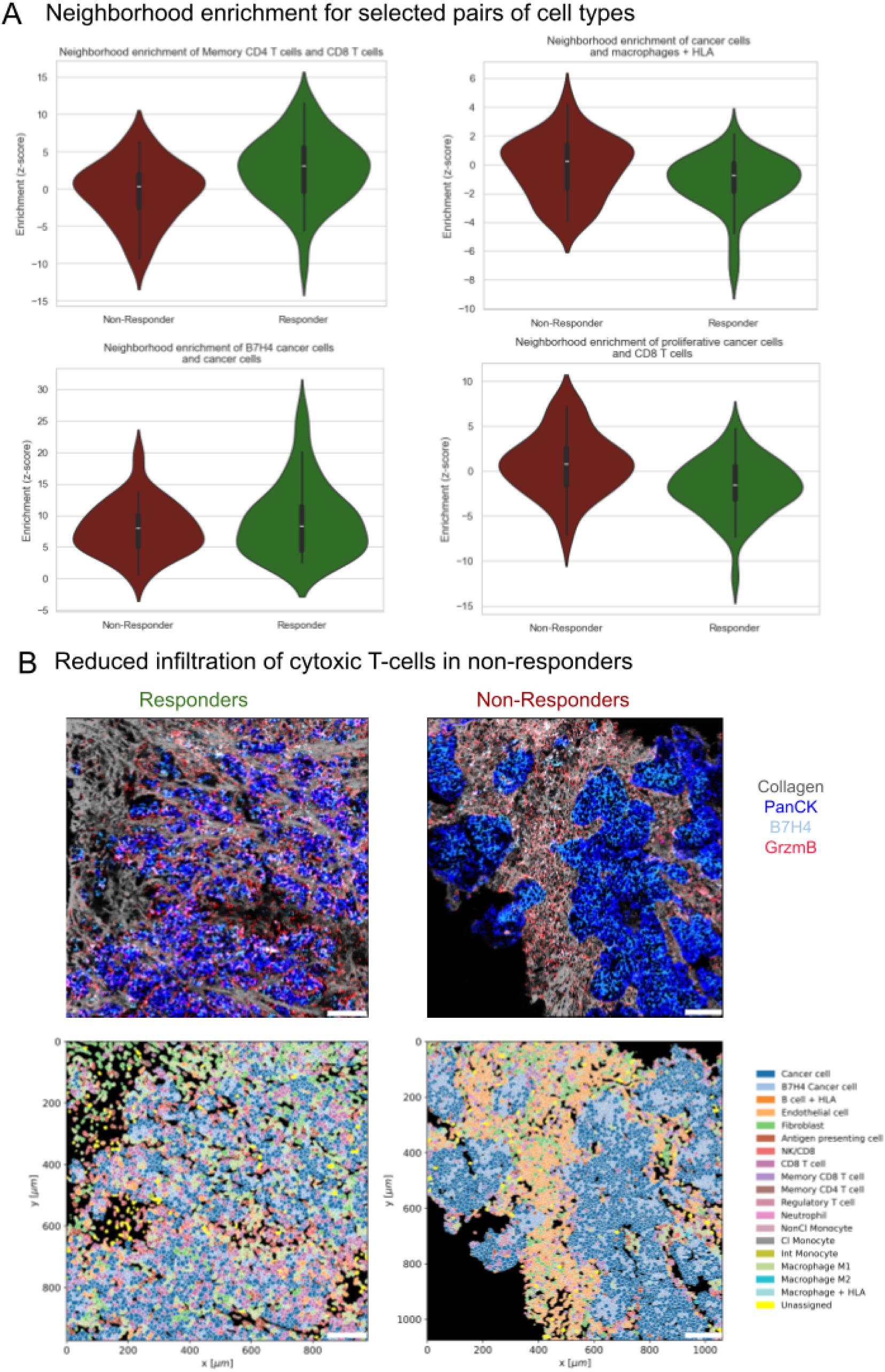
Spatial organization of cell types. **A** Distribution of neighborhood enrichment score over patients for selected pairs of cell types. The score quantifies proximity of a pair of cell types relative to random permutation of cells’ positions in an image, with positive values indicating co-localisation above random chance. Violins show the distribution across patients, and for each patient the neighbor enrichment was averaged across all ROIs of that patient. Mann-Whitney-U test statistics: CD4-CD8: 813 (p=0.005), Cancer-macrophage/HLA: 42 (p=0.06), B7H4-Cancer: 616 (p=0.67), Proliferative Cancer-CD8: 333 (p=0.003). **B** Representative raw and annotated images showing the colocalization of cancer cells (PanCK) and activated CD8+ T cells (GrzmB) in responders and non-responders. The scale bars are 125µm.

### CD8+ T cells are excluded from tumour regions in Non-Responders

Activated CD8 T expressing granzyme B (NK/CD8 cluster) exhibited no statistically significant differences in the average neighbourhood enrichment with any cancer cell cluster between responders and non-responders (Fig. S4-2B). However, this neighborhood enrichment score was calculated at the image level and averaged across patients in each response group which does not reflect local heterogeneity within the tissue. Indeed, individual samples exhibited regions of NK/CD8 cells (and other immune cells) infiltrating cancer cell groups in responder tissues, but not in responder samples where NK/CD8 cells were peripheral to the tumour and often separated by dense collagen (Fig. 4C). Protein abundance in the CD8/NK cell cluster further supported this with increased abundance of pan-keratin, β-catenin and E-cadherin in responder tissues, while collagen was more abundant in non-responder NK/CD8 cells (Fig. S4-2C). This is indicative of a fibrotic TME that could restrict immune cell infiltration, reducing anti-tumour immunity (Tharp et al. 2024; Mukund et al. 2025).

### Identifying spatial features that persist post-chemotherapy

To examine the trajectory of cell relationships in non-responder patient tissues before and after NACT, we performed similar analysis on matched tumour samples pre- and post-chemotherapy (Fig. 5A). The fraction of proliferating cancer cells over total cancer cells was slightly albeit significantly reduced in post-NACT samples; conversely, the proportion of proliferating B7H4 cancer cells was increased post-treatment, despite the overall proportion of B7H4+ cancer cells being reduced by chemotherapy(Fig. 5B), and no change in cell numbers overall (Fig. S5-1A). This is consistent with known increased efficacy of chemotherapy on proliferating cells, but indicates that B7H4 cancer cells may be more resistant to this.

**Figure 5.**
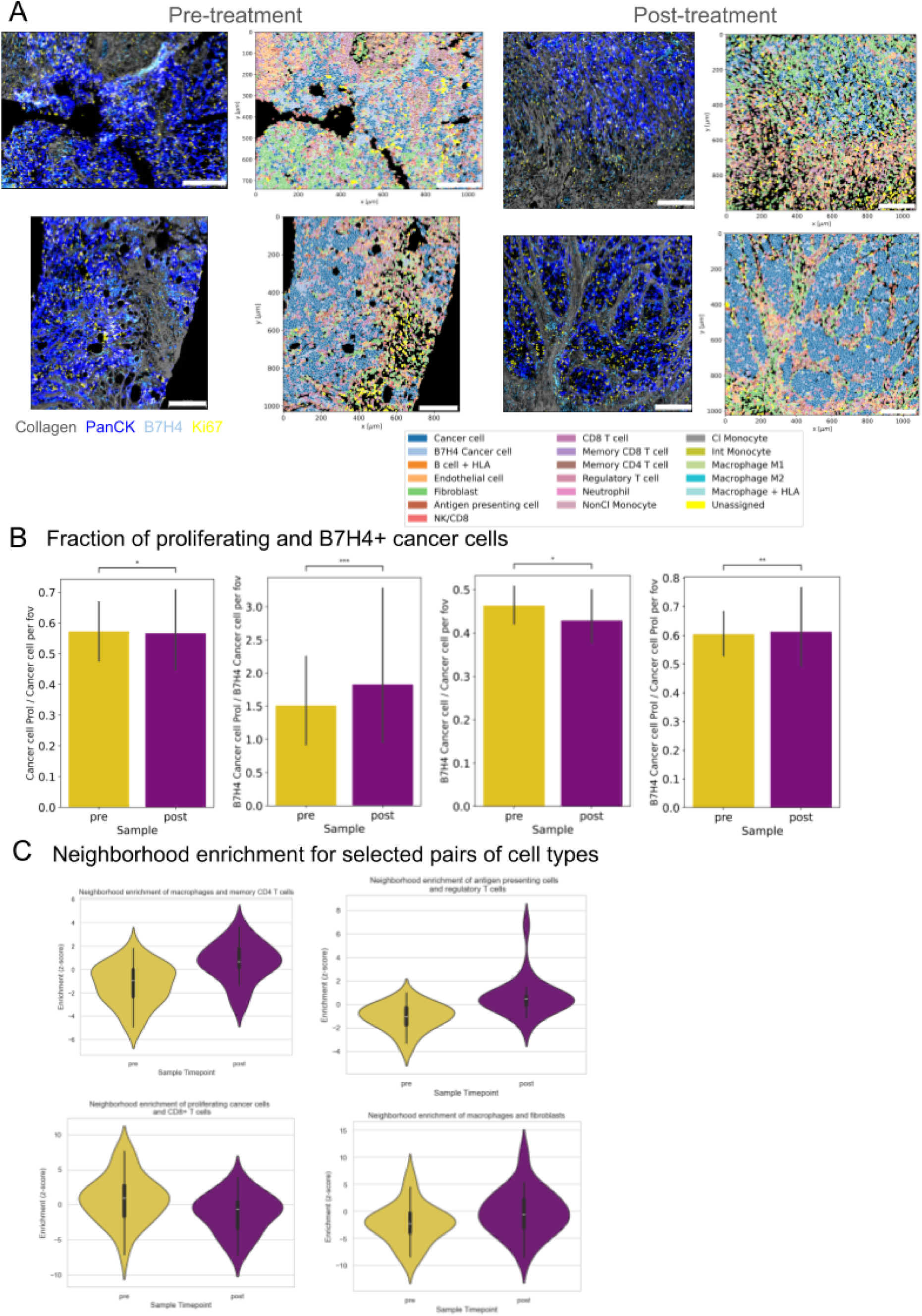
Tissue state before and after treatment. **A** Representative raw and annotated images from non-responder patients showing spatial organization of cell types (see legend). The pre- and post-treatment images are matched so that each row of images is from one patient. The scale bars are 200µm. **B** Fraction of cancer cells depending on B7H4 and proliferation (Ki67) expression. Bars show median and error bars show the standard error of the median. Stars represent the p-value for Kruskal-Wallis test as follows: *: 1.00e-02 < p <= 5.00e-02, **: 1.00e-03 < p <= 1.00e-02, ***: 1.00e-04 < p <= 1.00e-03. **C** Neighborhood enrichment in selected pairs of cell types. Violins show the distribution across patients, and for each patient the neighbor enrichment was averaged across all ROIs of that patient. Mann-Whitney-U test statistics: Macrophage-CD4: 117 (p=0.0006); APC-Treg: 81 (p=3e-5); Proliferative Cancer-CD8: 380 (p=0.05); Macrophage-fibroblast: 205, (p=0.1).

Next we analysed neighborhood enrichment to assess changes in spatial arrangement pre/post treatment. Macrophages and CD4 T cells were more co-localised post-treatment, as were antigen-presenting cells with regulatory T cells (Fig. 5C, top row). In contrast, proliferating cancer cell and CD8+ T cell colocalisation was reduced after treatment (Fig. 5C, bottom left). Macrophages and fibroblasts were more co-localised after treatment, indicating higher inflammation and fibrosis in post-treatment tissue (Fig. 5C, bottom right).

To further investigate features in non-responder patient samples that persist during chemotherapy as candidate ‘resistant’ signatures, we ranked pairs of cell types based on the absolute difference in their neighborhood enrichment, adding the differences between responders and non-responders pre-treatment (Fig. S4-2B) and between non-responders pre- and post-treatment (Fig.S5-1C). The top four pairs of cell types and their differences in average neighborhood enrichment are shown in Table 1, and the full distributions across patients in Fig. S5-2. Three of the four pairs of cell types were more co-localised in non-responders before treatment and even further enriched post-treatment, including fibroblasts with both M1 macrophages (Fig. S5-2B) and antigen presenting cells (Fig. S5-2C; Table 1 ranks 2&3), supporting previous reports of a fibroblast-macrophage interaction axis driving immunosuppression and suggesting this feature may be further exacerbated by NACT (Z. Liu et al. 2024). The enhanced co-localisation between blood vessels (endothelial cells) and Ki67-positive cancer cells was further increased post-treatment (Fig. S5-2D; Table 1, rank 4), indicative of establishment of a pro-metastatic niche. These findings demonstrate some features of non-responder tissues that persist and expand during NACT, and imply establishment of a stronger pro-invasive niche leading to tumour progression.

**Table 1.** Differences in neighborhood enrichment between responders and non-responders pre-treatment, and in non-responders between pre- and post-treatment samples. Pairs of cell types (second column) were ranked by the absolute value of their combined difference in neighborhood enrichment (fifth column), adding the difference in responders vs non-responders (third column), and pre- vs post-treatment (fourth column). A positive number means greater co-localisation in responders vs non-responders of pre- vs post-treatment, and a negative number means greater co-localisation in non-responders vs responders of post- vs pre-treatment.

| Rank | Cell Type Pairs | $\Delta NE R_{pre} - nR_{pre}$ | $\Delta NE nR_{pre} - nR_{post}$ | $\Delta NE Total$ |
| --- | --- | --- | --- | --- |
| 1 | B7H4 Cancer cell - B7H4 Cancer cell Prol | 1.20 | 2.81 | 4.01 |
| 2 | Macrophage M1 - Fibroblast | -1.73 | -2.04 | -3.76 |
| 3 | Antigen presenting cell - Fibroblast | -1.80 | -1.83 | -3.64 |
| 4 | Endothelial cell - Cancer cell Prol | -1.05 | -1.87 | -2.92 |

### Collagen analysis

Mechanical properties of tumour tissues change during growth and invasion, and recent evidence demonstrates that NACT can promote stiffening of TNBC tumours very early during treatment regimens, a feature that persists during the entire treatment period (Sinha et al. 2025). Examination of collagen can provide complementary information to such biomedical imaging techniques to define structural and mechanical contributions to spatial cell neighbourhoods. We measured 14 structural features of collagen (see Methods, with fibres identified via thresholding and edge detection (Figure 6A). Differences between responders and non-responders in pre-treatment samples were not statistically significant for any of the collagen features, though circular variance and fibre curvature were both higher in responders (Fig. S6-1A). Collagen fibres were less curved (Figure 6B, left) and more isotropic (Figure 6B, right) in post-treatment samples compared to pre-treatment, with no statistically significant differences observed in other collagen features.

**Figure 6.**
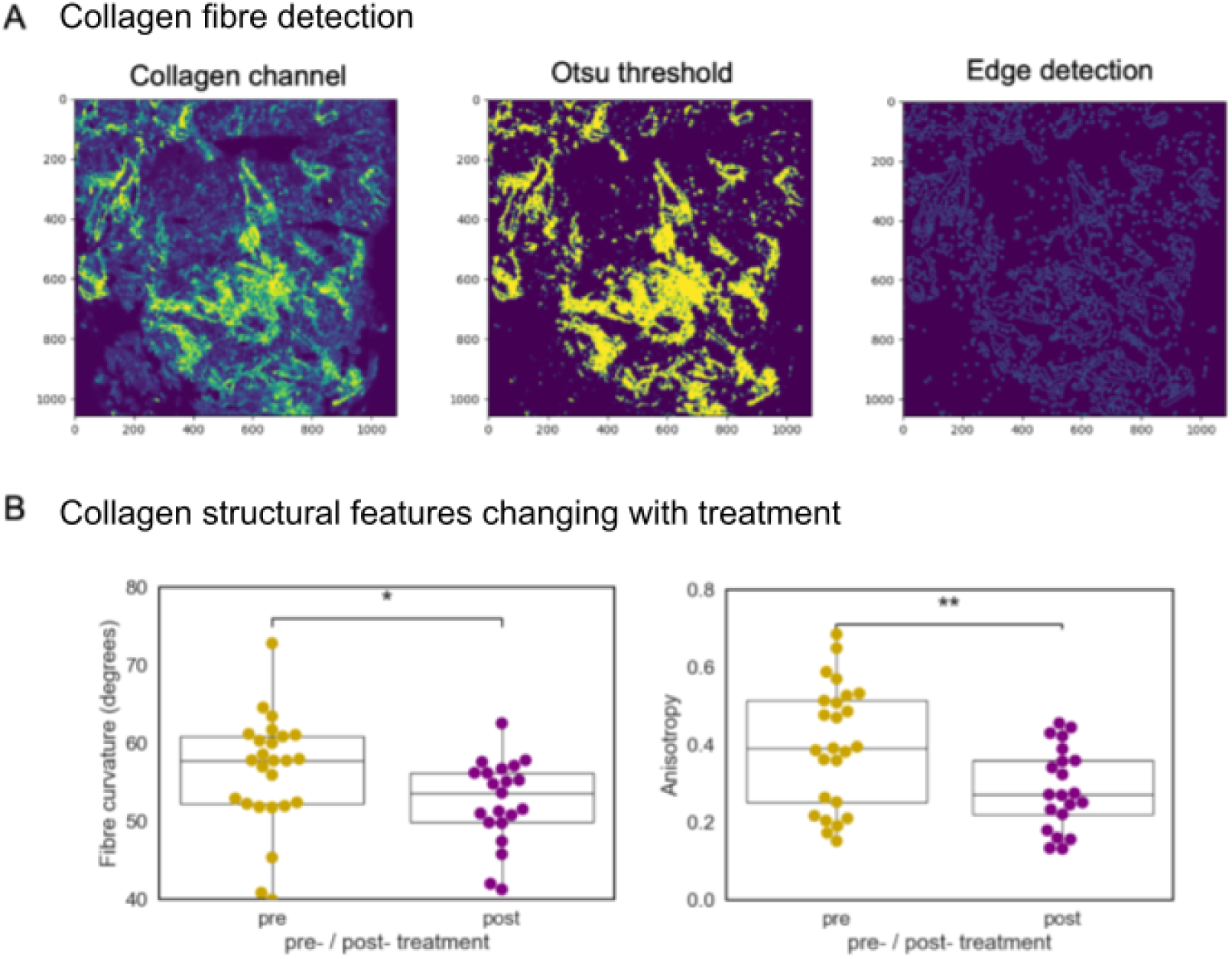
Collagen structure. **A** Representative example of segmentation of a collagen image. (right) original pre-processed collagen image, (middle) thresholded image (right) application of the Canny filter. **B** Comparison of distributions of collagen features pre- and post-treatment for non-responder patients only. (left) Collagen fibre curvature (t-stat = −2.03, p-value = 0.049), (right) Anisotropy (t-stat = −2.82, p-value = 0.007). Boxplots display median, upper and lower quartiles.

Given the complexity of these various spatial relationships and their relatively small individual effect sizes, we next sought to integrate multiple spatial and molecular features simultaneously using machine learning.

### GNN learns features from cell-cell contact graph and protein abundance that are predictive of chemotherapy response

While our analysis of cell type abundances and spatial arrangements revealed some differences between responders and non-responders in pre-treatment samples, these individual features showed relatively small effect sizes and high inter-patient variability. To comprehensively assess the predictive potential of the spatial-molecular data, we used a graph neural network (GNN) approach that can integrate multiple information layers simultaneously. GNNs are particularly well-suited for this task as they can capture complex relationships between spatially connected cells while incorporating the high-dimensional protein expression profiles that define each cell. In our implementation, each cell serves as a node in a spatial graph defined by cell-cell contacts, with additional feature vectors for each node consisting of either the 35-protein expression profile, cell type labels, or both.

Our GNN model achieved an AUROC of 0.71 for predicting chemotherapy response using pre-treatment data comprising protein abundance, cell type labels, and cell-cell contact graphs (Table 2). We evaluated three graph construction methods (k-NN, k-at-most, and Delaunay triangulation, see Methods) with different combinations of node features. The best performance was achieved using a k-at-most graph with combined protein abundance and cell type features, demonstrating that cell type information provides a small amount of additional predictive value beyond protein expression alone.

**Table 2:** Comparison of patient-level predictions of response to chemotherapy. GNN prediction accuracy (mean and std in AUROC scores, higher is better) using different graph construction and node features.

| Node Features | k-NN | K-Atmost | Delaunay |
| --- | --- | --- | --- |
| Protein Abundance | 0.68 +/- 0.07 | 0.68 +/- 0.07 | 0.68 +/- 0.07 |
| Cell-Type | 0.58 +/- 0.10 | 0.57 +/- 0.07 | 0.58 +/- 0.09 |
| Protein Abundance + Cell-Type | 0.70 +/- 0.08 | **0.71 +/- 0.07** | 0.70 +/- 0.07 |

To interpret the model’s predictions, we applied GNNExplainer (Ying et al. 2019) to identify the ten most informative features in each ROI (Figure 7A-B). To aggregate this information across ROIs from all patients, we counted how often each marker appeared in the top ten of any ROI, and ranked the markers (Figure 7C-D). The analysis revealed that **CD11b, CD366, FOXP3** and **B7H4** were the most predictive protein markers for treatment response across all ROIs, followed by **Pan-keratin, CD8a, Vimentin,** and **CD45RO**. We applied the same interpretability analysis to identify the most predictive cell types, which were **fibroblasts, cancer cells, and CD8 T cells**. These findings both recapitulate our spatial analysis results, and also revealed additional predictive features: The prominence of immune checkpoint molecules B7H4 and CD366 (TIM-3), as well as regulatory T cell marker FOXP3 indicates that immune regulatory states, rather than simple immune cell abundance, are contributing factors to chemotherapy response. The identification of CD45RO as a key predictor suggests that memory T-cell states may also influence treatment outcomes.

**Figure 7:**
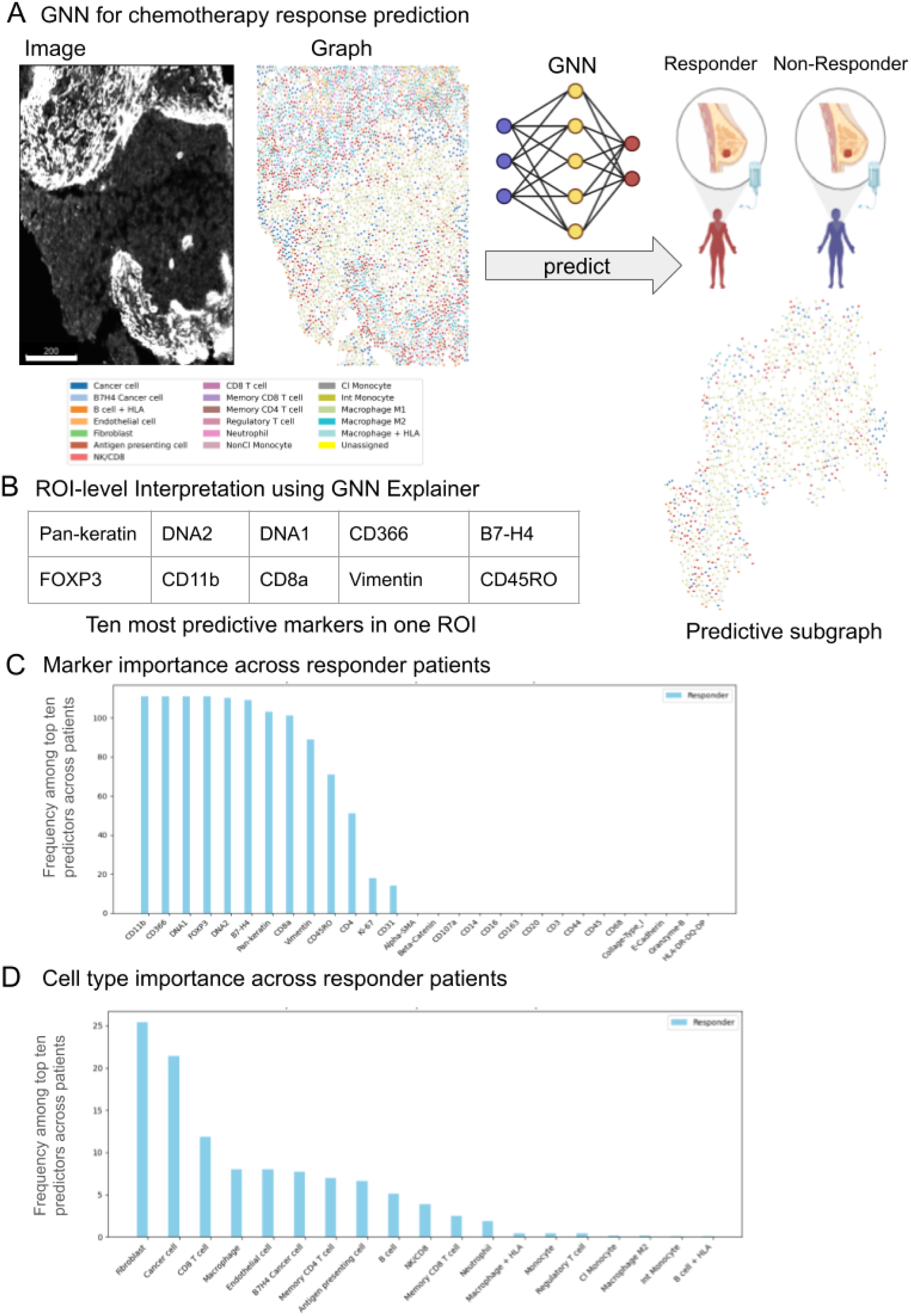
Response prediction and interpretability. **A** Graph neural network (GNN) model architecture integrating cell-cell contact graphs and protein abundance profiles to predict response to therapy. Each cell serves as a node with its 35-protein expression profile as features, connected by spatial proximity edges. Images on left show a representative IMC channel (scale bar is 200µm) and corresponding cell-cell contact graph, colored by cell type label (see legend). **B** Interpretability analysis for an example responder ROI using GNNExplainer yields a minimal predictive subgraph (right), and the ten protein markers contributing most to prediction performance (bottom). **C-D** Patient-level aggregation of protein marker importance scores (C) and cell type importance scores (D) associated with prediction of chemotherapy response, aggregated across all test samples from GNNExplainer analysis for each individual sample.

## Discussion

Our comprehensive spatial analysis of TNBC revealed several important findings: (1) individual cell type abundances showed only modest differences between chemotherapy responders and non-responders, with small effect sizes; (2) spatial relationships between cell types, particularly T-cell exclusion from tumor regions in non-responders, differed between responders and non-responders; (3) specific spatial features – including fibroblast-macrophage co-localization – persisted and intensified in non-responders during treatment, suggesting core resistance mechanisms; and (4) graph neural networks integrating spatial-molecular data achieved moderate prediction accuracy (AUROC=0.71), identifying immune regulatory markers as key predictive features. Together with the SMART study (Wall et al. 2025) our work illustrates how analysis of different modes of spatial data can reveal distinct determinants of therapeutic response.

### Spatial organization, not cell abundance, distinguishes treatment response

While we observed some differences in cell type frequencies between responders and non-responders—including increased antigen-presenting cells and regulatory T cells in responders—these differences had small effect sizes and substantial inter-patient variability. This finding aligns with emerging evidence that spatial context, rather than simple cell counts, drives therapeutic outcomes in cancer (Zheng et al. 2023; Punovuori et al. 2024; Bull et al. 2024; Mukund et al. 2025; X. Q. Wang et al. 2023; X. Wang et al. 2024). The limited predictive value of abundance data alone underscores why bulk analysis approaches may fail to predict treatment response in heterogeneous tumors like TNBC.

We identified CD8+ T cell exclusion from tumor regions in non-responders. This spatial exclusion pattern has been recognized as prognostic of response to adjuvant chemotherapy in TNBC (Loi et al. 2019). The mechanisms underlying this exclusion may include physical barriers such as collagen deposition or metabolic competition, though further investigation would be needed to confirm the specific drivers in our cohort.

While the SMART study (Wall et al. 2025) using spatial transcriptomics on the same patient cohort did not observe CD8+ T cell spatial exclusion patterns, it revealed seven distinct tumour-immune microenvironments (TIMEs) and identified B-cell enriched niches as prognostically significant—particularly those surrounding histologically normal epithelium. This highlights the complementary nature of our approaches: IMC’s single-cell resolution enables detection of fine-scale spatial exclusion patterns, while transcriptomics captures the molecular states underlying these spatial arrangements. Both studies converge on the importance of immune regulatory states, with Well et al.’s TIME classifications aligning with our identification of FOXP3 and CD366 as key predictive markers of immune regulation.

Our collagen analysis, while limited by the 2D nature of IMC and ROI selection constraints, showed trends toward straighter, more organized fibers in non-responders. This aligns with the extensive literature on tumor-associated collagen signatures (TACS), where organized collagen correlates with poor prognosis (Conklin et al. 2011).

### Persistent spatial features identify potential therapeutic targets

Our analysis of matched pre- and post-treatment samples from non-responders revealed spatial features that not only distinguish non-responders initially but persist or intensify during treatment. Fibroblast-macrophage and fibroblast-antigen presenting cell co-localisation are both already detectable pre-treatment in non-responders, suggesting that they could represent resistance mechanisms rather than secondary responses to chemotherapy toxicity. The persistent fibroblast-macrophage co-localization we observed suggests these interactions may represent a targetable resistance mechanism. Potential therapeutic approaches could include targeting macrophages through CSF1R inhibitors (Wen et al. 2023) or fibroblasts directly with fibroblast activation protein (FAP) ligands coupled to cytotoxic drugs (Sahai et al. 2020; Kim et al. 2017), or through targeting key mediators of their crosstalk, such as CSF and PDGF (Adler et al. 2020).

The increased co-localization between endothelial cells and proliferating cancer cells post-treatment hints at the establishment of a pro-metastatic vascular niche. This pattern aligns with evidence that chemotherapy can paradoxically enhance metastatic potential through vascular remodeling (Karagiannis et al. 2017), suggesting that vascular normalisation therapies might be particularly beneficial in TNBC patients showing this spatial signature.

While our analysis focuses on specific tumor regions, we acknowledge that spatial features may vary across the tumor microenvironment, which may impact the interpretation of persistent spatial features. While the annotation by pathologists is a strength of this dataset <u>(Wall et al. 2025)</u>, it also introduces bias in the selection of ROIs. Future studies can make use of whole-slide imaging methods to reduce that bias.

### Graph neural networks integrate spatial-molecular relationships and identify predictive features

Our GNN approach achieved moderate predictive accuracy (AUROC=0.71), successfully identifying biologically relevant features that individual spatial metrics missed. The prominence of B7H4, CD366, and FOXP3 as top predictive markers emphasizes the predictive capacity of immune regulatory states over simple immune cell presence.

Our use of GNNs contrasts Wang et al.’s (X. Q. Wang et al. 2023) approach, which used selected spatial features as input to logistic regression models to predict immunotherapy response with similar accuracy, but failed to predict response to chemotherapy alone from pre-treatment samples. We demonstrate that the more comprehensive use of features enabled by the GNN can extract predictive information for chemotherapy response specifically from spatial-molecular data in pre-treatment samples.

While our AUROC of 0.71 represents moderate accuracy, it demonstrates proof-of-principle that spatial-molecular features contain predictive information about chemotherapy response that could complement existing clinical markers.

### Computational pipeline contributions

Beyond the biological insights, our study provides a scalable computational framework for large-scale IMC analysis that addresses key technical challenges in spatial proteomics. Our end-to-end pipeline is assembled from existing state-of-the-art computational tools to provide a robust, modular and convenient foundation for future spatial proteomics studies. The implementation of image-level batch correction represents a relatively novel approach in IMC pipelines (Windhager et al. 2023), allowing for removal of batch effects prior to cell segmentation and cell type classification. However, we acknowledge the inherent risk of over-correction that could potentially remove genuine biological differences, and our approach is intended as a reasonable alternative to the methods imported from the single-cell transcriptomics field.

### Clinical implications and future directions

Our findings suggest potential refinements to current treatment strategies for TNBC patients. While chemotherapy combined with immunotherapy has been shown to improve therapy response and survival (Dixon-Douglas and Loi 2023), spatial features could inform which patients might benefit from the added immunotherapy or further targeted therapies. Patients showing persistent fibroblast-immune interactions during early treatment cycles might warrant early introduction of stromal-targeting agents. Future work should focus on combined analysis of multiple cohorts for greater statistical power, prospective validation, integration with existing clinical predictors, and development of clinically practical, cost-effective monitoring strategies based on these spatial signatures.

Fully-integrated analysis of multiple modalities in the SMART dataset (Wall et al. 2025) is a direction for future work, wherein end-to-end joint analysis can enable discovery in connected data beyond the comparison of single-modality results.

## Methods

### Data collection and preprocessing

We preprocessed IMC images to remove hot pixels and batch effects, followed by segmentation to extract protein profiles and locations of individual cells. We then clustered cells and labelled clusters according to their protein profiles. We compared cell type abundances as well as their spatial arrangement between different categories of patient samples. Finally, we use a GNN to predict whether patients will respond to chemotherapy and interpret what the predictions are based on. The complete framework for processing IMC data and predicting the response of patients to chemotherapy is in Figure 1.

### TNBC IMC Dataset

We use a recently generated IMC dataset, which is available on BioImage Archive under doi:10.6019/S-BIAD2027. The dataset consists of a combination of retrospective samples (via biobanking consent) and from the FORCE clinical trial (Sinha et al. 2025). All patients were diagnosed with triple negative breast cancer (TNBC) and had a tissue biopsy taken before starting neoadjuvant chemotherapy. FORCE trial patients also had biopsies taken after chemotherapy cycles 1.1 and 2.1. Patients received either EC-T (epirubicin, cyclophosphamide, taxanes) or EC-T with carboplatin (Fig. 1A) for neoadjuvant chemotherapy (NACT) regimens. Residual cancer burden (RCB) post-NACT treatment was determined (Fig. 1B, Fig. S1A). Samples were acquired from post-NACT surgical resection material from non-responder patients for analysis. Pathologist-guided regions of interest (ROIs) identified in sequential H&E tissue sections were then analyzed using IMC. IMC identified protein abundance through a panel of 35 metal-tagged antibody markers for tumoral, immune, and stromal cells (Fig. S2B; CD3, CD4, CD8a, CD11b, CD14, CD16, CD20, CD27, CD31, CD38, CD44, CD45, CD45RO, CD68, CD107a, CD163, CD366, Beta-Catenin, E-Cadherin, Pan-Keratin, Vimentin, Tbet, FOXP3, HLA-DR-DQ-DP, Alpha-SMA, Granzyme-B, B7-H4, Ki-67, PD1, PD-L1, PD-L2, p53, Collagen Type I, EGFR, VEGF) as well as two channels for DNA antibodies that are used for segmentation. This resulted in a pixel-resolved image with 35 channels (one for each protein marker). In total the dataset comprised 813 ROIs and over 4 million cells (Fig. 1B). We processed these images using a custom-assembled pipeline that includes image preprocessing, cell segmentation, cell phenotyping, spatial analysis, and chemotherapy response prediction (Fig. 1C, see Results and following Methods sections).

### IMC hot pixel detection and denoising

As IMC data contain both hot pixels and shot noise we preprocessed images to remove these. Hot pixels are thought to occur due to the formation of antibody aggregates (Lu et al. 2023) that result in a technical artefact of unusually high antibody counts. Shot noise arises because of the discrete nature of ion detection and antibody binding, which causes random fluctuations in signal intensity. These noise sources limit the precision with which we can determine protein expression levels, and we employ statistical methods to compensate for them: After extracting ROIs with Steinbock (Windhager et al. 2023) we process these with IMC-Denoise (Lu et al. 2023). IMC-Denoise identifies hot-pixel artefacts by evaluating their pixel neighbours and removing pixels that are much brighter than their neighbours, and furthermore uses a self-supervised deep-learning-based algorithm to correct for shot noise.

### Batch correction

We treat each protein channel as independent, and normalise its intensity into the interval [0,1] by using a contrast-limited adaptive histogram equalisation (CLAHE) algorithm (Pizer et al. 1987)., which splits the image into regions and equalises the distribution of pixel intensities towards a more uniform distribution. Adaptive histogram equalisation improves the contrast of the image, and in the context of IMC, it can help to correct for antibodies with different binding affinities. In CLAHE the maximum contrast transformation is limited, to avoid over-amplifying background noise from regions of low contrast.

### Cell segmentation

After image-level batch correction we apply the Mesmer method (Greenwald et al. 2022), a segmentation algorithm to delineate individual cells and quantify protein expression levels within distinct cellular regions. Mesmer is a deep learning-based algorithm that provides cell masks for spatial localization of proteins. It takes both cytoplasmic and nuclear channels as input, and segments both nuclei and whole cells as output. For our choice of channels for segmentation see Fig. S2B. Once the cells are segmented, centroids are computed to represent the cells’ spatial coordinates within an ROI. The segmented cells are used for cell phenotyping and to generate a single cell proteomic view that is used for cell type labelling.

To mitigate the impact of cell segmentation errors, we check for the following quality control criteria: 1) Cells without nucleus (2.6% of the cells) may appear because we are analysing 2D slices of a 3D structure, but also occur due to segmentation artefacts. Thus, we decided to remove these from further analysis. 2) Cells with a nucleus to cytoplasm size ratio (N/C) > 1 (2.8% of the cells) were caused because nuclei were assigned to the wrong cells and/or multiple cells (Fig. S2C). Since we are only using the whole cell segmentation mask (but not the nuclear mask) for cell type annotation and the downstream analysis, this misassignment of the nuclei doesn’t affect our analysis, and therefore we decided to keep those cells.

### Cellular phenotyping

To annotate the cells, we use the Pixie method (C. C. Liu et al. 2023), a pipeline which uses a two-step clustering process to create cell-level features based on clustering pixels from segmented cells, and then cell clusters based on these features. For the clustering we include only phenotypic markers which allow us to identify cell types, while functional markers that can be expressed in various cell types, such as Ki-67, are excluded (Fig. 3a). We excluded IMC channels where antibody staining was judged to be of insufficient quality (CD27, CD38, T-bet, p53, EGFR, VEGF, PD-1, PD-L1, PD-L2). Given a set of image channels (the user-chosen markers), Pixie first clusters pixels using a self-organising map, and then assigns these clusters into meta-clusters using consensus hierarchical clustering. If necessary, the final clusters can be manually refined and annotated with biologically relevant labels. After pixel clusters have been identified and annotated, the frequency of each pixel cluster within each cell is used as the feature vector for cell clustering. Cells are clustered using the same process of self-organising maps followed by hierarchical clustering and manual refinement. To label cell clusters as biologically relevant cell types and states, we made an initial annotation (Fig. S3-1A) of the cells using the table shown in Figure S3-1C. We then inspected and adjusted the clusters that were not well separated during cell clustering as follows: We defined manual thresholds based on marker intensity to combine or separate specific clusters (Fig. S3-1B). For those cases in which it was not obvious where to set the threshold based on bimodality of the intensity distribution, visual inspection of the images was carried out to assess the marker positivity. This method is also useful to inspect those markers that were not directly used for phenotyping, but whose expression is important to identify individual cell states, such as the proliferative state, based on the marker Ki-67. A detailed list of thresholds and the clusters annotated is in Table S1 and figure S3-1B.

The combination of markers present in one of the cell clusters made them difficult to annotate, likely due to “lateral spillover” between neighbouring cells, as well as some unrealistically large cell segmentation masks. We therefore labeled them as “Unassigned”. We checked our final annotation by comparing the protein abundance in each cell compared to all other cells (Fig. S3-2).

### Cell type abundances and spatial arrangement

After cell annotation we analyse the cell type abundance and their spatial arrangement. Frequency of cell-cell contacts was quantified using the neighborhood enrichment function in SquidPy (Palla et al. 2022). We chose to conduct a hypothesis-driven examination of selected cell pairs over a comprehensive exploratory analysis. Therefore, we did not test all pairs of cell types for statistical significance in the difference of neighborhood enrichment, and as such did not use multiple-testing correction in Figs. 4A and 5C.

### Collagen feature analysis

An Otsu threshold was applied to CLAHE-processed collagen images to determine regions of collagen fibres. Collagen features were obtained using scikit-image for Python (van der Walt et al. 2014) and The Workflow of Matrix Biology Informatics (TWOMBLI) (Wershof et al. 2021) workflow for ImageJ software. Collagen area was determined as the fraction of the image occupied by fibre pixels. Lacunarity, a descriptor of spatial heterogeneity of fibres organisation, was assessed by convolution of a 30 × 30 pixel kernel to compute local fibre mass sums across each image then calculate their distribution. Collagen compactness was calculated as the average distance from a pixel to the background for all fibre regions. Fibre perimeter and solidity were derived via scikit-image’s ‘regionprops’ functions. Anisotropy of fibres within an image was calculated from the eigenvalues of the inertia tensor of an image, determined using ‘regionprops’. Anisotropy was taken to be the ratio of secondary and primary eigenvalues subtracted from unity. Individual fibres were extracted by applying a Canny edge detector filter to the image. Extracted fibres were used to determine average fibre length and circular variance of fibres. Local fibre orientations were computed from Sobel gradients of fibre edges. Circular variance was calculated as 1 − |𝑧|, where |𝑧| is the mean complex orientation vector. Lower circular variance indicates greater fibre alignment. The proportion of fibre branchpoints and endpoints per unit area were calculated using TWOMBLI. TWOMBLI was also used to measure the box-counting fractal dimension of an image, alignment of fibres and fibre curvature, measured as the change in angle along a collagen fibre over a baseline distance (40 micrometres).

### Prediction of response to chemotherapy

We use a graph neural network (GNN) model to predict whether patients will respond to chemotherapy. We only use patients with EC-T and EC-T-carboplatin chemotherapy treatments. Patients are classified based on their residual cancer burden (RCB) score at the end of the treatment. Here, we adopt a binary label for the patient response, defining a pathological, complete responder (pCR) if the RCB is 0 and non-responder otherwise (nR). We remove ROIs with less than 1000 cells. The final dataset used for prediction consists of 63 patients with a total of 636 ROIs. Among these patients, 37 are responders and 26 are non-responders. For developing our ML model, we split the patient cohort into training and validation sets using a 70-30 ratio, resulting in 45 patients (446 ROIs) for training and 18 patients (190 ROIs) for evaluation. We run the model with five different random splits and report mean and standard deviation in AUROC score.

### Graph construction

The cell-cell contact graph provides insight into the spatial connectivity of cells at varying scales, and thus captures potential cellular interactions. In our work, we combine this graph-based representation with protein expression data. We test three distinct graph construction approaches: k-nearest neighbours (kNN), k-almost neighbours, and Delaunay triangulation. The kNN method connects each cell to its k nearest neighbours based on the Euclidean distance between centroids. In contrast, the k-atmost neighbour approach uses a distance threshold. It connects each cell with up to k cells if their distance falls within the specified threshold while ensuring each cell is at least connected to one nearest neighbour. The threshold here is the weight that separates the weakest 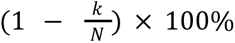 of edges (where N is the number of nodes in a graph). Finally, the Delaunay triangulation method connects the centroids of cells forming triangles such that no cell centroid lies inside the circumcircle of any triangle, inferring cellular adjacency based on their spatial arrangement. We construct graphs with each of these methods and investigate their performance on the downstream task of predicting chemotherapy response.

### GNN architecture

Let 𝑋 ∈ 𝑅*^n×d^* be a protein expression matrix, 𝐺 = (𝑉, 𝐸) be cell-cell contact graph with 𝑛 = |𝑉| as a set of vertices and |𝐸| as a set of edges, 𝑑 is the number of protein markers, and 𝑦 ∈ {0, 1} be a label that takes value 1 for responder or 0 for non-responder. We use a GNN encoder 𝑓_θ_ to embed input 𝐺 and 𝑋 of each ROI to a fixed length representation 𝑧 ∈ 𝑅^d𝑧^, where 𝑑_𝑧_ is the dimensionality of latent space, 𝑔_ϕ_ to predict responder/non-responder labels. The encoder and response predictor parameters jointly optimise binary cross-entropy loss *L*

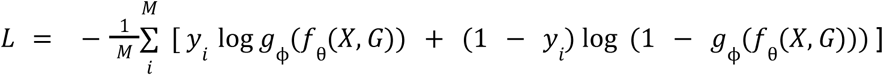

where θ and ϕ are the parameters of the encoder and response predictor, respectively. The final loss is averaged over all 𝑀 ROIs in the dataset. The loss is optimised using a stochastic optimizer Adam (Kingma and Ba 2014) with a learning rate of10^−3^. The encoder architecture comprises a two-layer Graph Convolutional Network (GCN) (Kipf and Welling 2017) followed by average pooling. The response prediction head is a linear layer that predicts the probability of a patient’s response to chemotherapy.

### ROI to patient-level prediction

In the present dataset each patient has multiple ROIs. We first treat each ROI as separate samples for training purposes, before predicting a patient-level response which is more relevant for any clinical decision-making. During the evaluation phase, patient-level predictions were obtained using a majority voting scheme across the ROIs belonging to the same patient. Specifically, let 𝑅_𝑝_ = {𝑟_1_, 𝑟_2_, …, 𝑟_𝑛_} represent the set of ROI-level predictions for patient 𝑝, where 𝑟𝑖 ∈ {0, 1} represents the predicted response (0 for non-responder, 1 for responder). The patient-level prediction 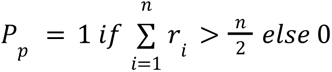.

### Model development, training and evaluation

We implemented our GNN framework in Python using PyTorch (Paszke et al. 2017) and PyTorch Geometric library (Fey and Lenssen 2019) for GNNs. To evaluate the predictive performance of the model, we report the area under the curve (AUC), accuracy, and F1 score to capture the effect of class imbalance. We report these scores for evaluation of ROI-level prediction and patient-level prediction using majority voting.

We split our dataset into a training and test set with a 70% and 30% split. We set k=7 for the k-NN and k-atmost graph construction. We run all experiments with 10 random seeds and report the mean and standard deviation. We experimented with different k values from the set {1, 2, 3, 4, 5, 6, 7, 8, 9, 10} and found that k=7 performed best.

### Interpretability of GNN prediction results

To interpret the predictions of the GNN-based model, we used GNNExplainer (Ying et al. 2019) to identify features critical for predicting chemotherapy response. GNNExplainer assigns importance scores to protein markers and edges within the cell-cell contact graph, reflecting their contributions to the model’s predictive performance. Using these scores, we retained the top 15% of edges in the cell-cell contact graph and identified the top 10 protein markers that consistently contributed to the model’s predictions across the test set.

### Code availability

Python code for all analysis can be found in the following github repositories: Preprocessing, denoising, and segmentation: https://github.com/Schumacher-group/IMC_preprocessing

Cell annotation and spatial analysis: https://github.com/Schumacher-group/IMC_TNBC_analysis

GNN Prediction: https://github.com/Schumacher-group/ML4SpatialAnalysis

## Supporting information

all figures incl. supplementary figures

## Acknowledgements

LL, AK, and GT thank Leor Rose for helpful discussions.

## Rights Retention Statement

For the purpose of open access, the author has applied a Creative Commons Attribution (CC BY) licence to any Author Accepted Manuscript version arising from this submission.

## Potential conflicts of interest

CS is on the scientific advisory board of Cytoreason Ltd.

---

**Table S1.** Thresholds used for refinement of initial cell clusters.

| Marker | Unrefined cluster | Threshold (t) | new cluster (< t) | new cluster (> t) |
| --- | --- | --- | --- | --- |
| CD20 | CD8 T cell | 0.98 | CD8 T cell | B cell |
| CD20 | Antigen presenting cell | 0.98 | Antigen presenting cell | B cell |
| B7-H4 | Cancer cell | 0.70 | Cancer cell | B7-H4 cancer cell |
| E-Cadherin | Antigen presenting cell | 0.30 | Antigen presenting cell | Cancer cell |
| FoxP3 | Memory CD4 T cell | 0.96 | Memory CD4 T cell | Regulatory T cell |
| FoxP3 | B7-H4 cancer cell | 0.96 | B7-H4 cancer cell | Regulatory T cell |
| CD31 | Fibroblast | 0.94 | Fibroblast | Endothelial cell |
| Granzyme B | Neutrophil | 0.94 | Neutrophil | NK/CD8 |
| CD163 | Macrophage | 0.90 | Macrophage | Macrophage M2 |
| CD107a | Macrophage M2 | 0.90 | Macrophage M2 | NK/CD8 |
| E-Cadherin | Memory CD8 T cell | 0.40 | Memory CD8 T cell | Cancer cell |
| CD14 | Monocyte | 0.60 | NonCI Monocyte | CI Monocyte |
| CD16 | CI Monocyte | 0.64 | CI Monocyte | Int Monocyte |
| HLA | Macrophage | 0.86 | Macrophage M1 | Macrophage + HLA |
| HLA | Macrophage M2 | 0.86 | Macrophage M2 | Macrophage + HLA |
| Ki-67 | Cancer cell | 0.40 | Cancer cell | Cancer cell Prol |
| Ki-67 | B7-H4 cancer cell | 0.40 | B7-H4 cancer cell | B7-H4 cancer cell Prol |

## Notes

### Competing Interest Statement

Potential conflict of interest: C.S is on the scientific advisory board of Cytoreason Ltd.

### Summary of Updates

In this version we have included supplementary figures that were omitted from the initial automated upload from eLife.

### Funding

- Wellcome Leap

