## Supplementary material for "Identifying tissue states by spatial protein patterns related to chemotherapy response in triple-negative breast cancer": all figures incl. supplementary figures

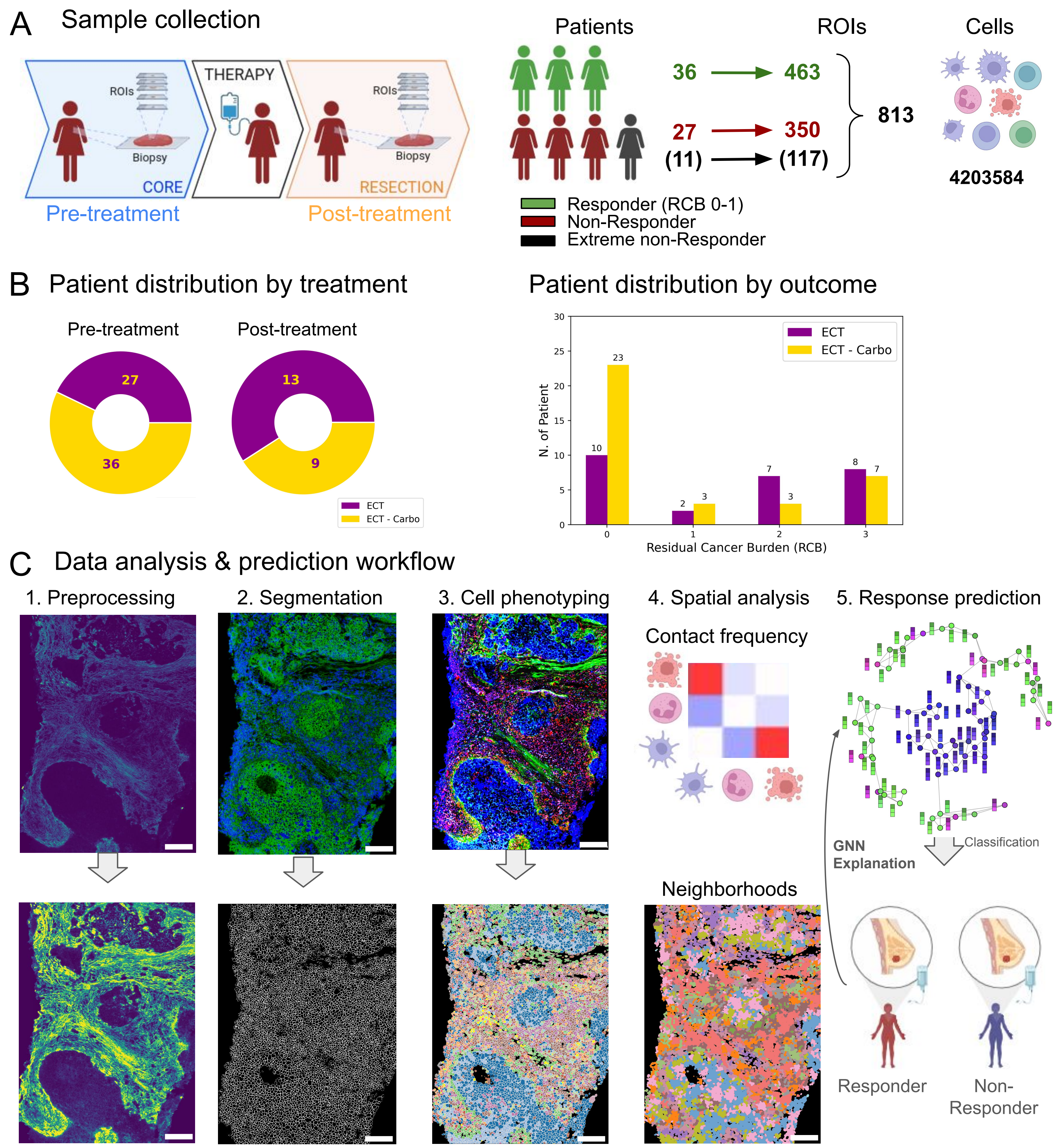


**Figure 1 - Overview of dataset and analysis workflow. A** Biopsies were taken before and after NACT (left), and multiple regions of interest (ROIs) extracted from each biopsy, stained with metal-tagged antibodies and imaged using Imaging Mass Cytometry (IMC). Number of ROIs in each response class (middle) and total number of cells identified in the dataset (right). **B** All patients received EC-T chemotherapy, and some patients with carboplatin (EC-T-carbo (left). Distribution of patients over residual cancer burden (right). **C.** Analysis workflow for IMC data showing an illustrative example of preprocessing, segmentation, cell phenotype, spatial analysis, and response prediction. The scale bar is 150μm. Image colours show the following: 1. Preprocessing: intensity for representative IMC channel. 2. Segmentation: Representative IMC channels in green and blue (top) and segmentation mask (bottom). 3. Cell phenotyping: Various IMC channels used for cell phenotyping (top) and cell type labels after segmentation (bottom). 4. Spatial analysis: schematic heatmap illustrating high (red) or low (blue) co-localisation between pairs of cell types (represented by cartoon drawings) (top) and cells coloured by neighborhood cluster label (bottom). 5. Response prediction: cell are nodes on the graph, edges represent contact between nearest neighbors, and each cell also carries a feature vector (here represented by columns) of protein expression and/or cell type label.

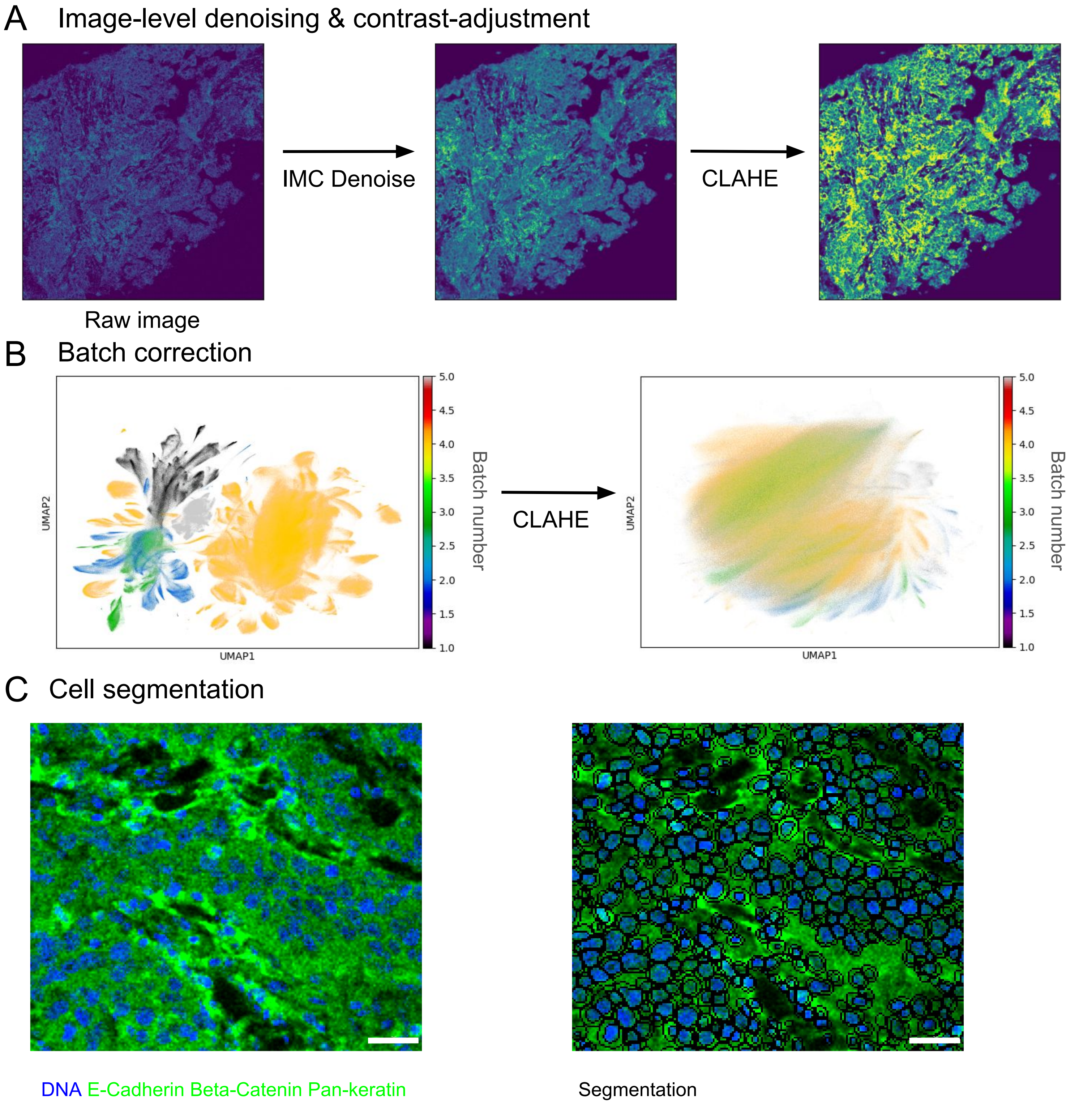


**Figure 2 - Data preparation: denoising, batch effects, segmentation.** **A** Denoised and contrast enhanced image of a representative ROI. **B** Approximate overlap of data from different patients reflects removal of batch effects. Each point is a single cell. After histogram equalisation (CLAHE) the batches are no longer separated. **C** Representative raw image (left) and with overlaid segmentation boundaries for nuclei and whole cells (right). Blue: DNA antibody; Green: Combination of E-Cadherin, Beta-Catenin, and Pan-keratin staining. The scale bar is 25µm

A

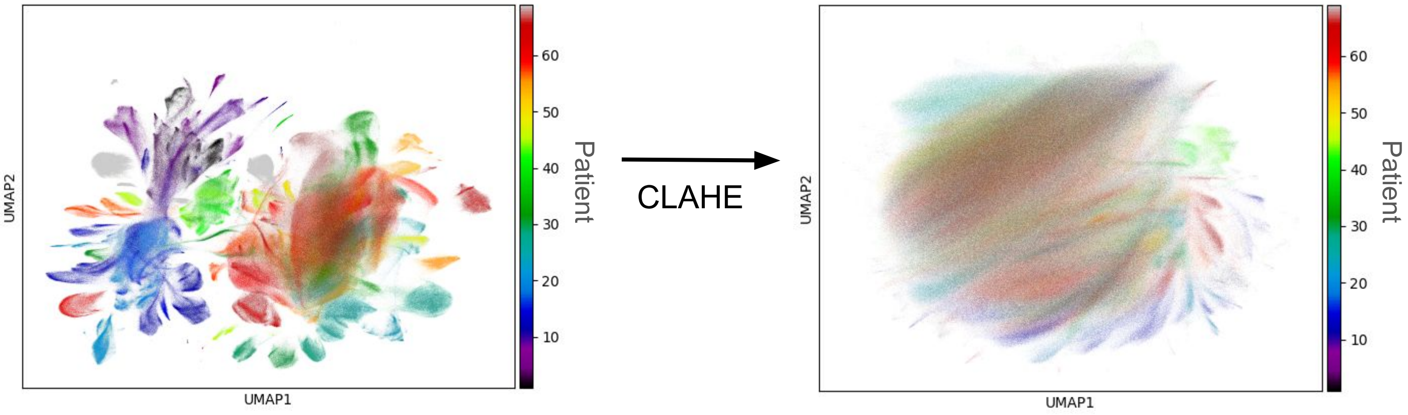


B

| Channels | | | |
|---|---|---|---|
| Alpha-SMA | CD27 | Collagen Type 1 | PD-1 |
| B7-H4 | CD31 | DNA1/DNA2 | PD-L1 |
| Beta-Catenin | CD38 | E-Cadherin | PD-L2 |
| CD3 | CD44 | EGFR | Tbet |
| CD4 | CD45 | FoxP3 | VEGF |
| CD8a | CD45RO | Granzyme-B | Vimentin |
| CD11b | CD68 | HLA-DR-DQ-DP | cytoplasm |
| CD14 | CD107a | Ki-67 | nuclei |
| CD16 | CD163 | p53 | state |
| CD20 | CD366 | Pan-Keratin | phenotype |

C

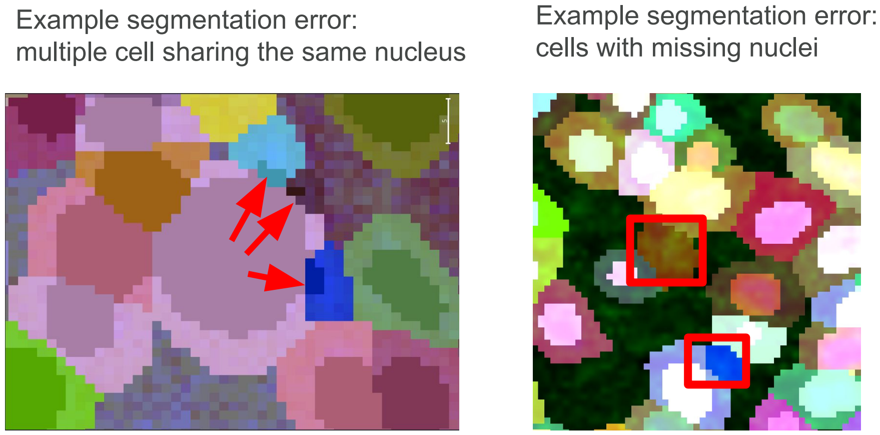


**Figure S2 (supplementary figure related to Figure 2) A** Approximate overlap of data from different patients reflects removal of batch effects. After histogram equalisation (CLAHE) the batches are no longer separated. **B** List of IMC channels, of which green highlighted one were used for cell segmentation, and DNA1/DNA2 for nuclear segmentation. **C** Representative example of segmentation errors that can occur. The scale bar is 5µm.

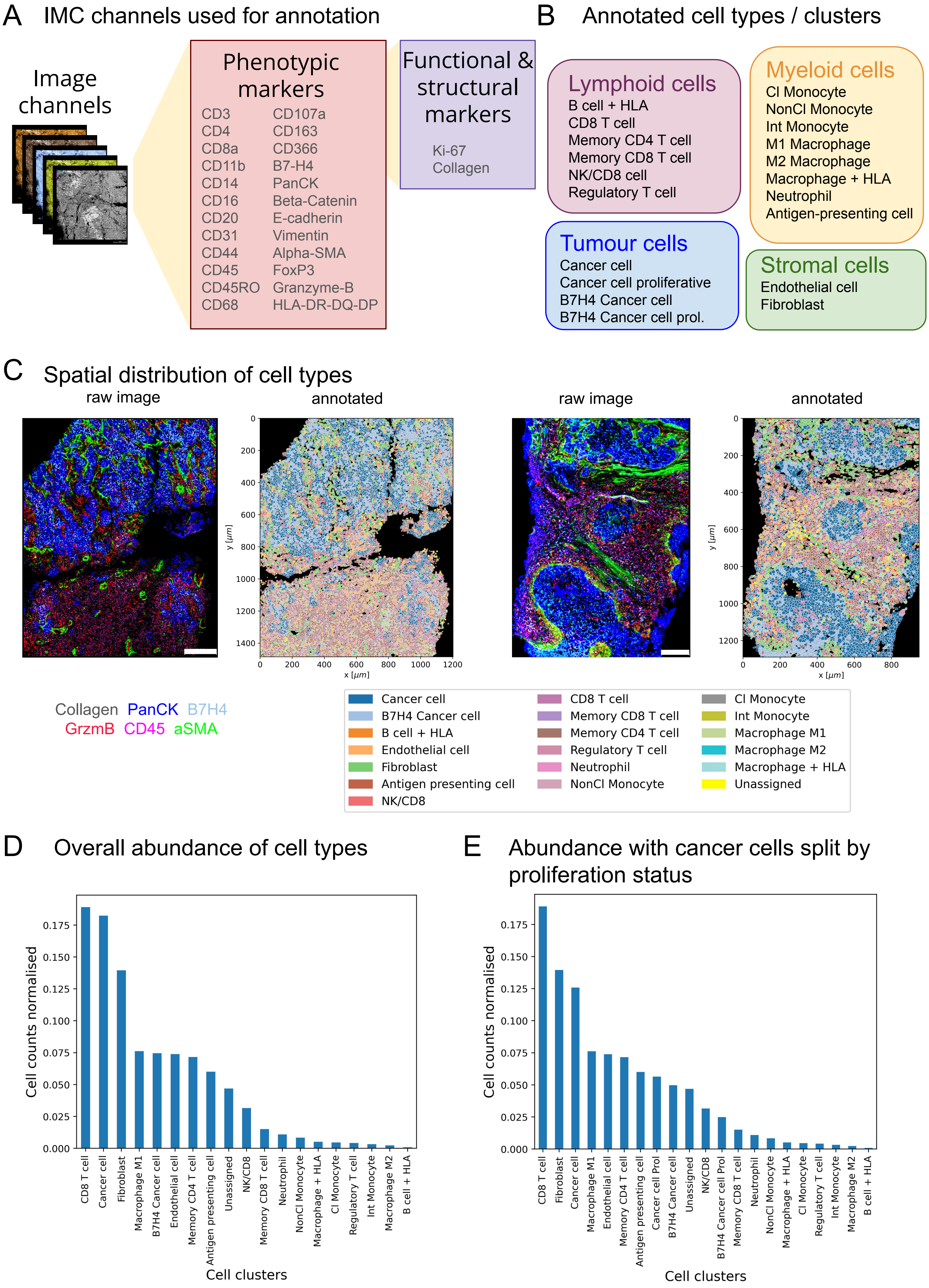
A IMC channels used for annotation
Image channels
Phenotypic markers
CD3 CD4 CD8a CD11b CD14 CD16 CD20 CD31 CD44 CD45 CD45RO CD68
CD107a CD163 CD366 B7-H4 PanCK Beta-Catenin E-cadherin Vimentin Alpha-SMA FoxP3 Granzyme-B HLA-DR-DQ-DP
Functional & structural markers
Ki-67 Collagen
B Annotated cell types / clusters
Lymphoid cells
B cell + HLA
CD8 T cell
Memory CD4 T cell
Memory CD8 T cell
NK/CD8 cell
Regulatory T cell
Myeloid cells
Cl Monocyte
NonCl Monocyte
Int Monocyte
M1 Macrophage
M2 Macrophage
Macrophage + HLA
Neutrophil
Antigen-presenting cell
Tumour cells
Cancer cell
Cancer cell proliferative
B7H4 Cancer cell
B7H4 Cancer cell prol.
Stromal cells
Endothelial cell
Fibroblast
C Spatial distribution of cell types
raw image
annotated
raw image
annotated
y [μm]
x [μm]
Collagen PanCK B7H4
GrzmB CD45 aSMA
Cancer cell
B7H4 Cancer cell
B cell + HLA
Endothelial cell
Fibroblast
Antigen presenting cell
NK/CD8
CD8 T cell
Memory CD8 T cell
Memory CD4 T cell
Regulatory T cell
Neutrophil
NonCl Monocyte
Cl Monocyte
Int Monocyte
Macrophage M1
Macrophage M2
Macrophage + HLA
Unassigned
D Overall abundance of cell types
Cell counts normalised
Cell clusters
CD8 T cell, Cancer cell, Fibroblast, Macrophage M1, B7H4 Cancer cell, Endothelial cell, Memory CD4 T cell, Antigen presenting cell, Unassigned, NK/CD8, Memory CD8 T cell, Neutrophil, NonCl Monocyte, Macrophage + HLA, Cl Monocyte, Regulatory T cell, Int Monocyte, Macrophage M2, B cell + HLA
E Abundance with cancer cells split by proliferation status
Cell counts normalised
Cell clusters
CD8 T cell, Fibroblast, Cancer cell, Macrophage M1, Endothelial cell, Memory CD4 T cell, Antigen presenting cell, Cancer cell Prol, B7H4 Cancer cell, Unassigned, NK/CD8, B7H4 Cancer cell Prol, Memory CD8 T cell, Neutrophil, NonCl Monocyte, Macrophage + HLA, Cl Monocyte, Regulatory T cell, Int Monocyte, Macrophage M2, B cell + HLA

**Figure 3 - Cell type assignment and abundance.** **A** IMC channels used for annotation of cell type (phenotypic markers) and functional state (functional markers). **B** Cell clusters annotated. **C** Representative raw images of selected channels (Collage, PanCK, B7H4, GrzmB, CD45, αSMA) and corresponding cell type labels (see legend). The scale bars are 200µm on the left and 150µm on the right. **D** Relative abundance of cell clusters across all ROIs, without subdividing proliferating cells. **E** Relative abundance of cell clusters across all ROIs, with sub-divided Cancer cell and B7H4 Cancer cell clusters into proliferation and non-proliferating based on Ki67 expression.

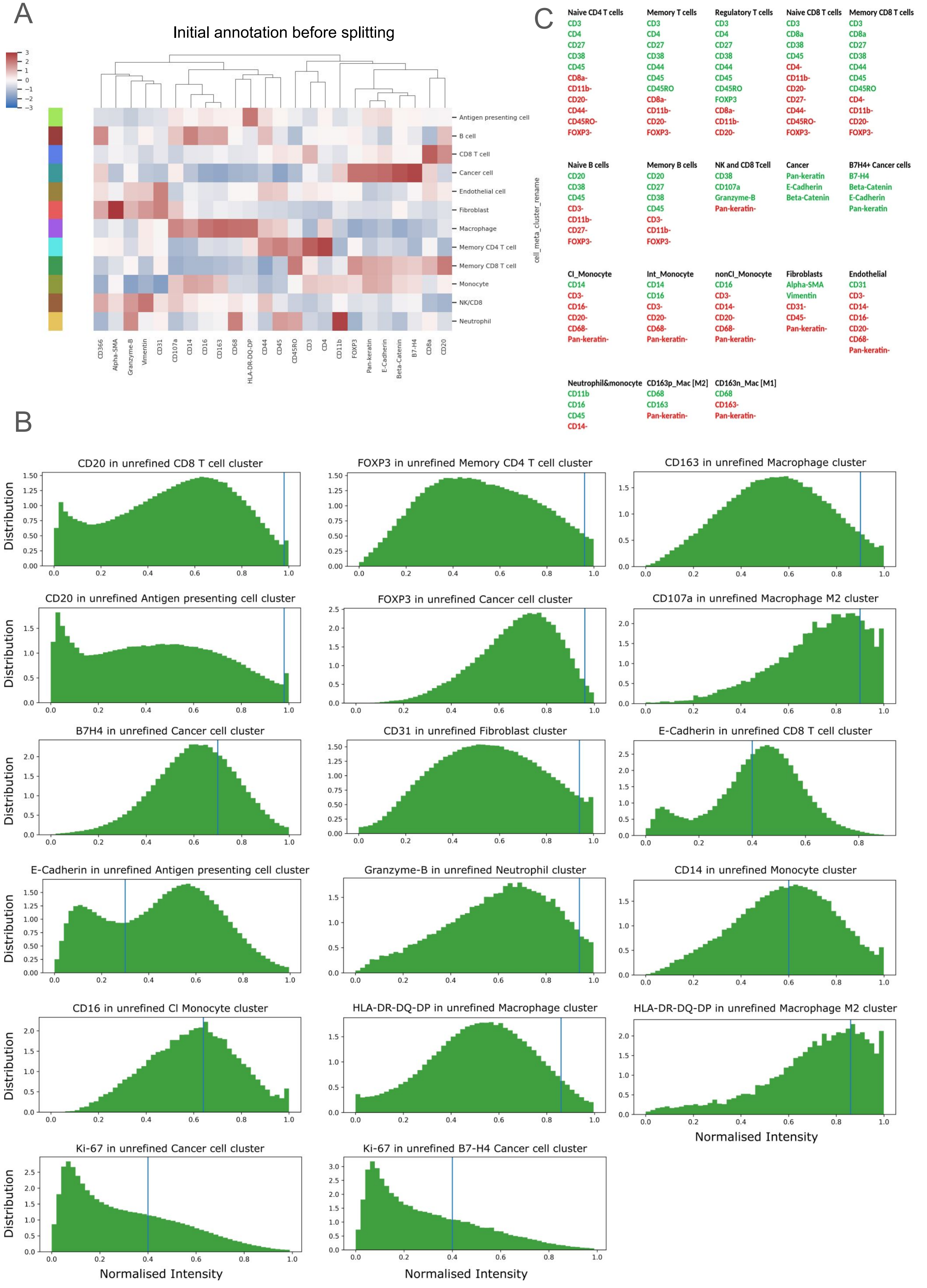
A
Initial annotation before splitting
Antigen presenting cell
B cell
CD8 T cell
Cancer cell
Endothelial cell
Fibroblast
Macrophage
Memory CD4 T cell
Memory CD8 T cell
Monocyte
NK/CD8
Neutrophil
cell_meta_cluster_rename
CD366
Alpha-SMA
Granzyme-B
Vimentin
CD31
CD107a
CD14
CD16
CD163
CD68
HLA-DR-DQ-DP
CD44
CD45
CD45RO
CD3
CD4
CD11b
FOXP3
Pan-keratin
E-Cadherin
Beta-Catenin
B7-H4
CD8a
CD20
C
Naive CD4 T cells
CD3
CD4
CD27
CD38
CD45
CD8a-
CD11b-
CD20-
CD44-
CD45RO-
FOXP3-
Memory T cells
CD3
CD4
CD27
CD38
CD44
CD45
CD45RO
CD8a-
CD11b-
CD20-
FOXP3-
Regulatory T cells
CD3
CD4
CD27
CD38
CD44
CD45
CD45RO
FOXP3
CD8a-
CD11b-
CD20-
Naive CD8 T cells
CD3
CD8a
CD38
CD45
CD4-
CD11b-
CD20-
CD27-
CD44-
CD45RO-
FOXP3-
Memory CD8 T cells
CD3
CD8a
CD27
CD38
CD44
CD45
CD45RO
CD4-
CD11b-
CD20-
FOXP3-
Naive B cells
CD20
CD38
CD45
CD3-
CD11b-
CD27-
FOXP3-
Memory B cells
CD20
CD27
CD38
CD45
CD3-
CD11b-
FOXP3-
NK and CD8 Tcell
CD38
CD107a
Granzyme-B
Pan-keratin-
Cancer
Pan-keratin
E-Cadherin
Beta-Catenin
B7H4+ Cancer cells
B7-H4
Beta-Catenin
E-Cadherin
Pan-keratin
Cl_Monocyte
CD14
CD3-
CD16-
CD20-
CD68-
Pan-keratin-
Int_Monocyte
CD14
CD16
CD3-
CD20-
CD68-
Pan-keratin-
nonCl_Monocyte
CD16
CD3-
CD14-
CD20-
CD68-
Pan-keratin-
Fibroblasts
Alpha-SMA
Vimentin
CD31-
CD45-
Pan-keratin-
Endothelial
CD31
CD3-
CD14-
CD16-
CD20-
CD68-
Pan-keratin-
Neutrophil&monocyte
CD11b
CD16
CD45
CD14-
CD163p_Mac [M2]
CD68
CD163
Pan-keratin-
CD163n_Mac [M1]
CD68
CD163-
Pan-keratin-
B
CD20 in unrefined CD8 T cell cluster
FOXP3 in unrefined Memory CD4 T cell cluster
CD163 in unrefined Macrophage cluster
CD20 in unrefined Antigen presenting cell cluster
FOXP3 in unrefined Cancer cell cluster
CD107a in unrefined Macrophage M2 cluster
B7H4 in unrefined Cancer cell cluster
CD31 in unrefined Fibroblast cluster
E-Cadherin in unrefined CD8 T cell cluster
E-Cadherin in unrefined Antigen presenting cell cluster
Granzyme-B in unrefined Neutrophil cluster
CD14 in unrefined Monocyte cluster
CD16 in unrefined Cl Monocyte cluster
HLA-DR-DQ-DP in unrefined Macrophage cluster
HLA-DR-DQ-DP in unrefined Macrophage M2 cluster
Ki-67 in unrefined Cancer cell cluster
Ki-67 in unrefined B7-H4 Cancer cell cluster
Distribution
Normalised Intensity

**Figure S3-1 (supplementary figure related to Figure 3) A** Initial unrefined cell types from SOM clustering of pixel clusters (see Methods) before refining cell types by splitting cluster based on channels shown in B. **B** Histograms of normalised intensity of relevant IMC channels in cell types that we refined by splitting. **C** List of cell types and which IMC channels were expected to be high (green) or low/absent (red), which was used for splitting of cell clusters and annotation.

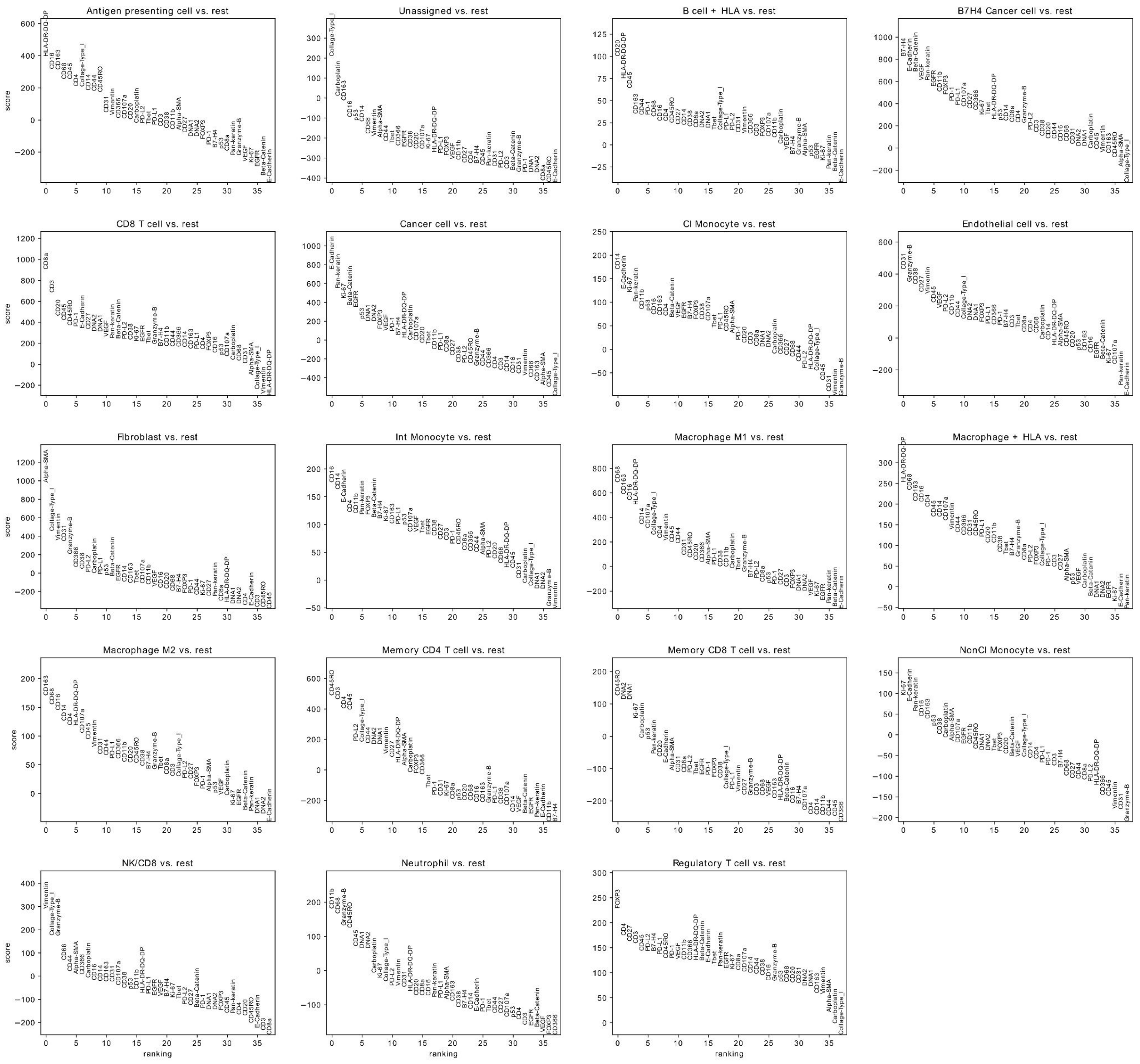


**Figure S3-2 (supplementary figure related to Figure 3)** Ranked wilcoxon scores of IMC channels in each cell type compared to all other cells.

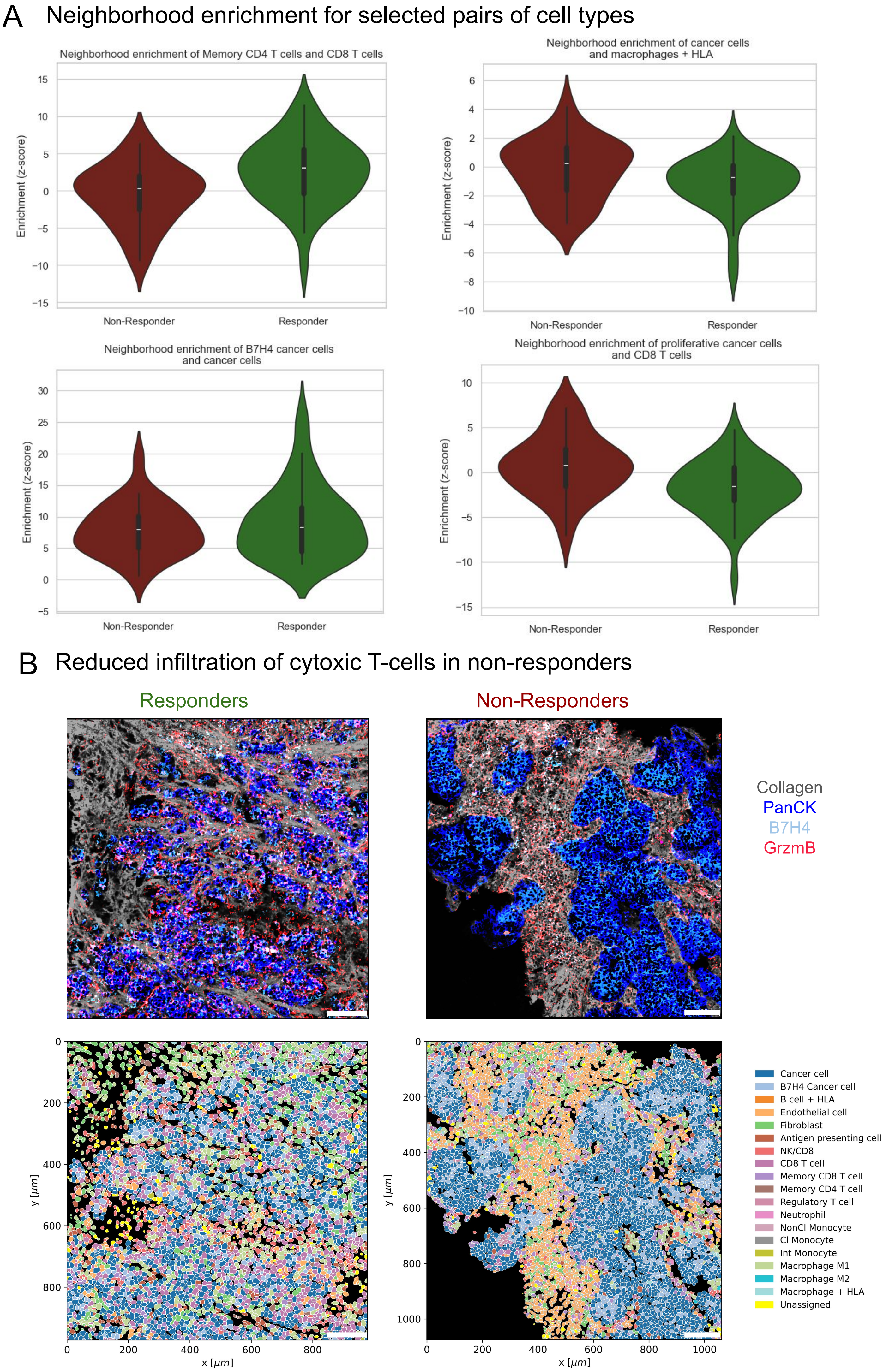
A
Neighborhood enrichment for selected pairs of cell types
Neighborhood enrichment of Memory CD4 T cells and CD8 T cells
Neighborhood enrichment of cancer cells and macrophages + HLA
Neighborhood enrichment of B7H4 cancer cells and cancer cells
Neighborhood enrichment of proliferative cancer cells and CD8 T cells
Enrichment (z-score)
Non-Responder
Responder
B
Reduced infiltration of cytoxic T-cells in non-responders
Responders
Non-Responders
Collagen
PanCK
B7H4
GrzmB
x [μm]
y [μm]
Cancer cell
B7H4 Cancer cell
B cell + HLA
Endothelial cell
Fibroblast
Antigen presenting cell
NK/CD8
CD8 T cell
Memory CD8 T cell
Memory CD4 T cell
Regulatory T cell
Neutrophil
NonCl Monocyte
Cl Monocyte
Int Monocyte
Macrophage M1
Macrophage M2
Macrophage + HLA
Unassigned

**Figure 4 - Spatial organization of cell types. A** Distribution of neighborhood enrichment score over patients for selected pairs of cell types. The score quantifies proximity of a pair of cell types relative to random permutation of cells' positions in an image. Violins show the distribution across patients, and for each patient the neighbor enrichment was averaged across all ROIs of that patient. Mann-Whitney-U test statistics: CD4-CD8: 813 (p=0.005), Cancer-macrophage/HLA: 42 (p=0.06), B7H4-Cancer: 616 (p=0.67), Proliferative Cancer-CD8: 333 (p=0.003). **B** Representative raw and annotated images showing the colocalization of cancer cells (PanCK) and activated CD8+ T cells (GrzmB) in responders and non-responders. The scale bars are 125µm.

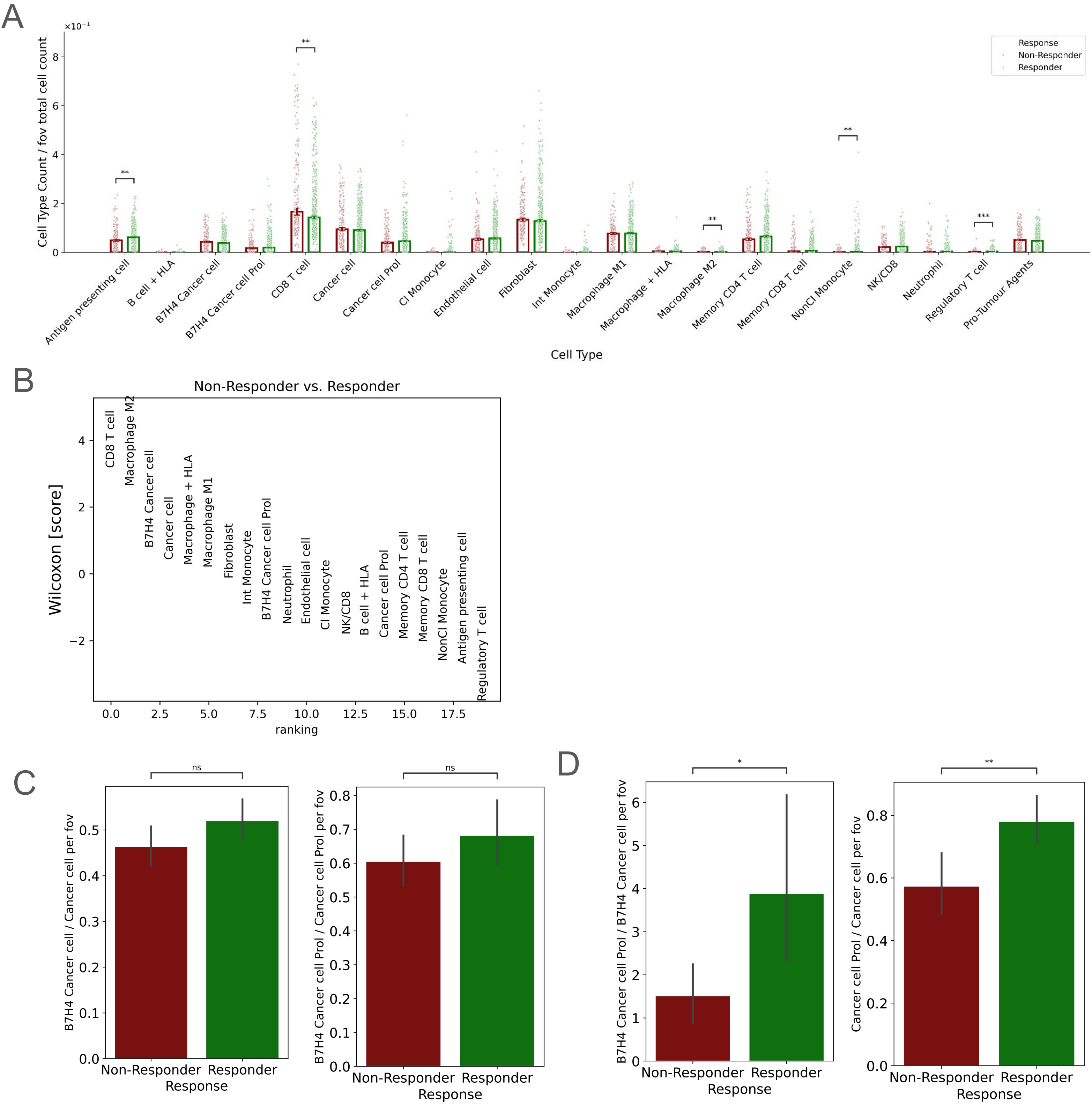


**Figure S4-1 (supplementary figure related to Figure 4) A** Number of each cell type relative to total number of cells per field of view (fov) for responders (green) and non-responders (red). **B** Wilcoxon rank-sum test score of the number of cells of each type shown in A in responders compared to non-responders. **C** Number of B7H4 Cancer cells relative to all Cancer cells (left) and proliferating B7H4 Cancer cells relative to all proliferating cancer cells. ns: not statistically significant (Kruskal-Wallis test). **D**. Number of proliferating B7H4 Cancer cells relative to all B7H4 Cancer cells (left) and proliferating Cancer cells relative to all Cancer cells.

Bars show median and error bars show the standard error of the median. Stars represent the p-value for Kruskal-Wallis test as follows: ns: 5.00e-02 < p <= 1.00e+00, *: 1.00e-02 < p <= 5.00e-02, **: 1.00e-03 < p <= 1.00e-02, ***: 1.00e-04 < p <= 1.00e-03, and we used the Benjamini-Hochberg correction for multiple testing in A.

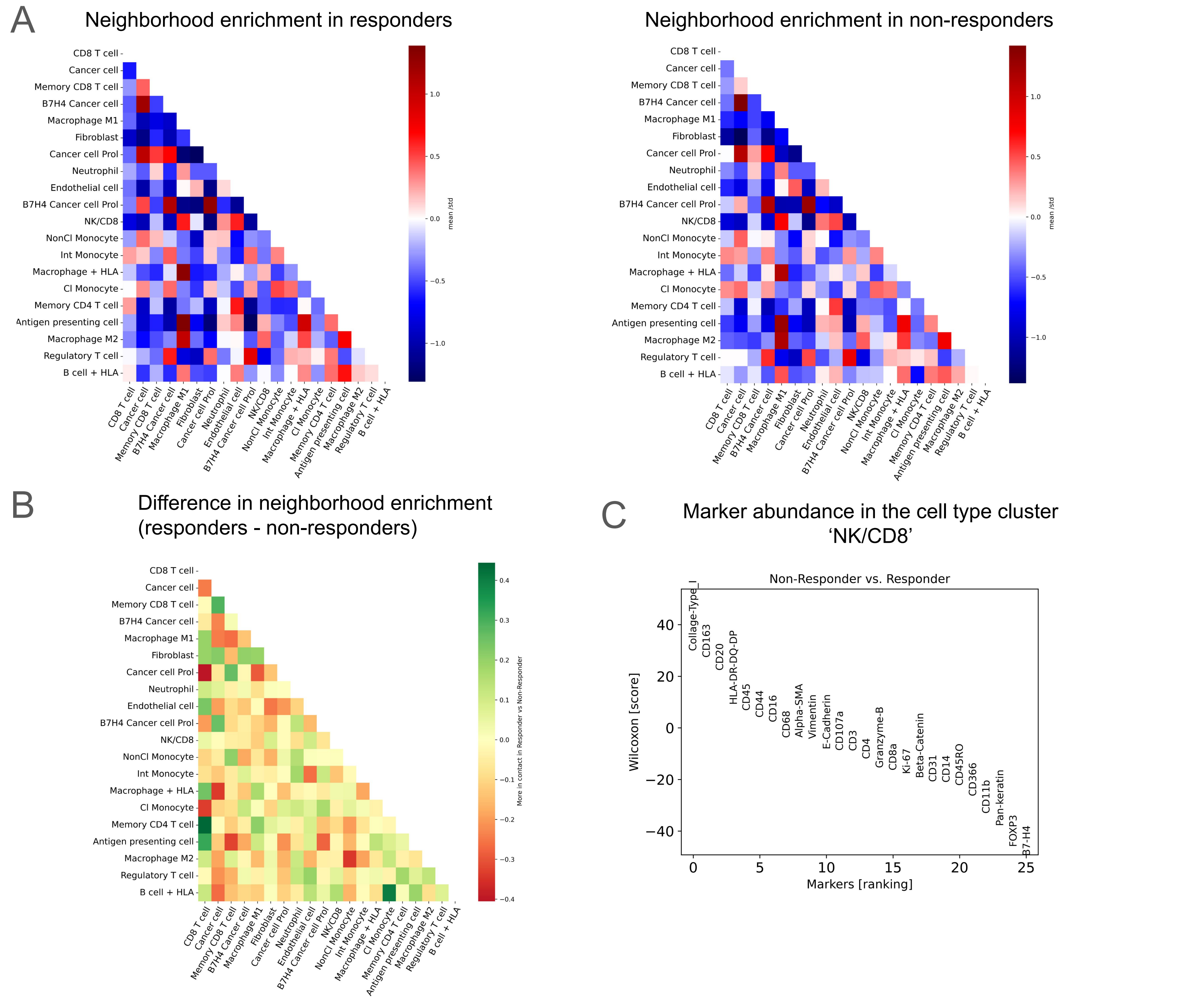


**Figure S4-2 (supplementary figure related to Figure 4) A** Neighborhood enrichment between all pairs of cell types in responders (left) and non-responders (right), averaged across all ROIs. **B** Difference of neighborhood enrichment in responders and non-responders **C** Marker abundance in the cell type cluster 'NK/CD8'.

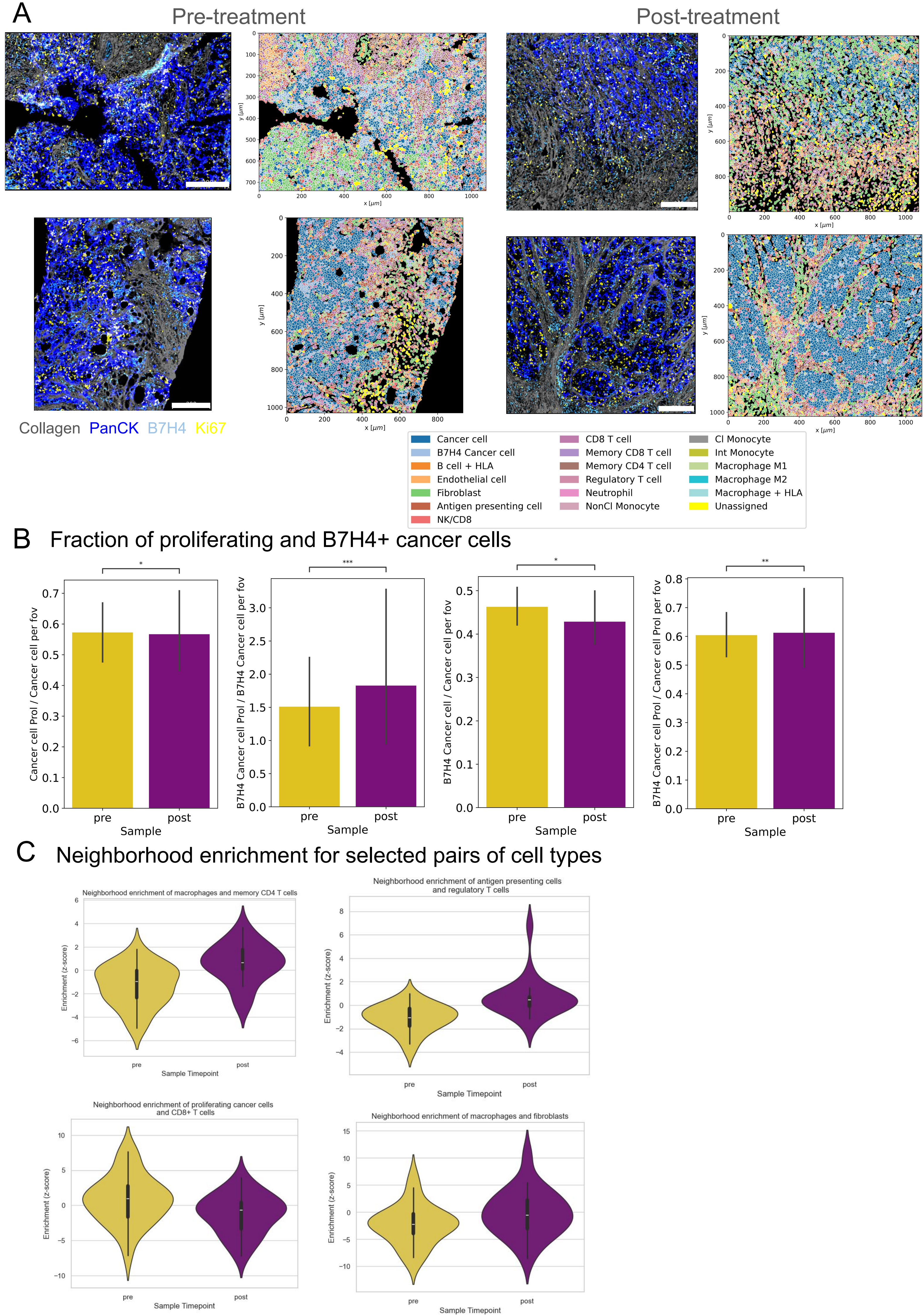
A
Pre-treatment
Post-treatment
Collagen PanCK B7H4 Ki67
x [μm]
y [μm]
Cancer cell
B7H4 Cancer cell
B cell + HLA
Endothelial cell
Fibroblast
Antigen presenting cell
NK/CD8
CD8 T cell
Memory CD8 T cell
Memory CD4 T cell
Regulatory T cell
Neutrophil
NonCl Monocyte
Cl Monocyte
Int Monocyte
Macrophage M1
Macrophage M2
Macrophage + HLA
Unassigned
B
Fraction of proliferating and B7H4+ cancer cells
Cancer cell Prol / Cancer cell per fov
B7H4 Cancer cell Prol / B7H4 Cancer cell per fov
B7H4 Cancer cell / Cancer cell per fov
B7H4 Cancer cell Prol / Cancer cell Prol per fov
*
***
*
**
pre
post
Sample
C
Neighborhood enrichment for selected pairs of cell types
Neighborhood enrichment of macrophages and memory CD4 T cells
Neighborhood enrichment of antigen presenting cells and regulatory T cells
Neighborhood enrichment of proliferating cancer cells and CD8+ T cells
Neighborhood enrichment of macrophages and fibroblasts
Enrichment (z-score)
Sample Timepoint

**Figure 5 - Tissue state before and after treatment. A** Representative raw and annotated images from non-responder patients showing spatial organization of cell types (see legend). The pre- and post-treatment images are matched so that each row of images is from one patient. The scale bars are 200µm. **B** Fraction of cancer cells depending on B7H4 and proliferation (Ki67) expression. Bars show median and error bars show the standard error of the median. Stars represent the p-value for Kruskal-Wallis test as follows: *: 1.00e-02 < p <= 5.00e-02, **: 1.00e-03 < p <= 1.00e-02, ***: 1.00e-04 < p <= 1.00e-03. **C** Neighborhood enrichment in selected pairs of cell types. Violins show the distribution across patients, and for each patient the neighbor enrichment was averaged across all ROIs of that patient. Mann-Whitney-U test statistics: Macrophage-CD4: 117 (p=0.0006); APC-Treg: 81 (p=3e-5); Proliferative Cancer-CD8:380 (p=0.05); Macrophage-fibroblast: 205, (p=0.1).

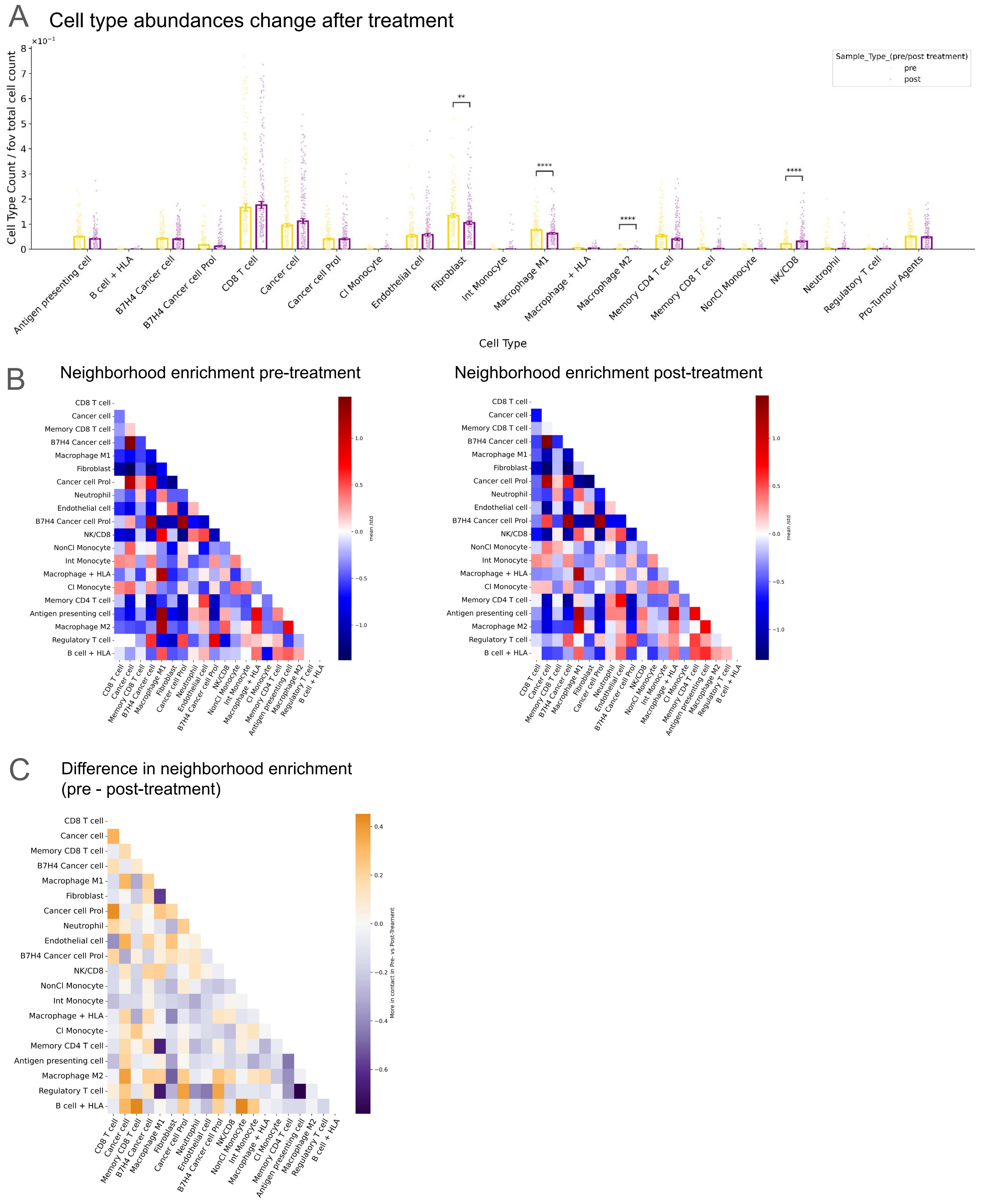


**Figure S5-1 (supplementary figure related to Figure 5) A** Number of each cell type relative to total number of cells per field of view (fov) for pre-treatment samples (yellow) and post-treatment samples (purple). Bars show median and error bars show the standard error of the median. Stars represent the p-value for Kruskal-Wallis test as follows: **: 1.00e-03 < p <= 1.00e-02,  ***: 1.00e-04 < p <= 1.00e-03,  ****: p <= 1.00e-04, and we used the Benjamini-Hochberg correction for multiple testing. **B** Neighborhood enrichment between all pairs of cell types in pre-treatment (left) and post-treatment (right) samples, averaged across all ROIs. **C** Difference of neighborhood enrichment in pre- and post-treatment samples.

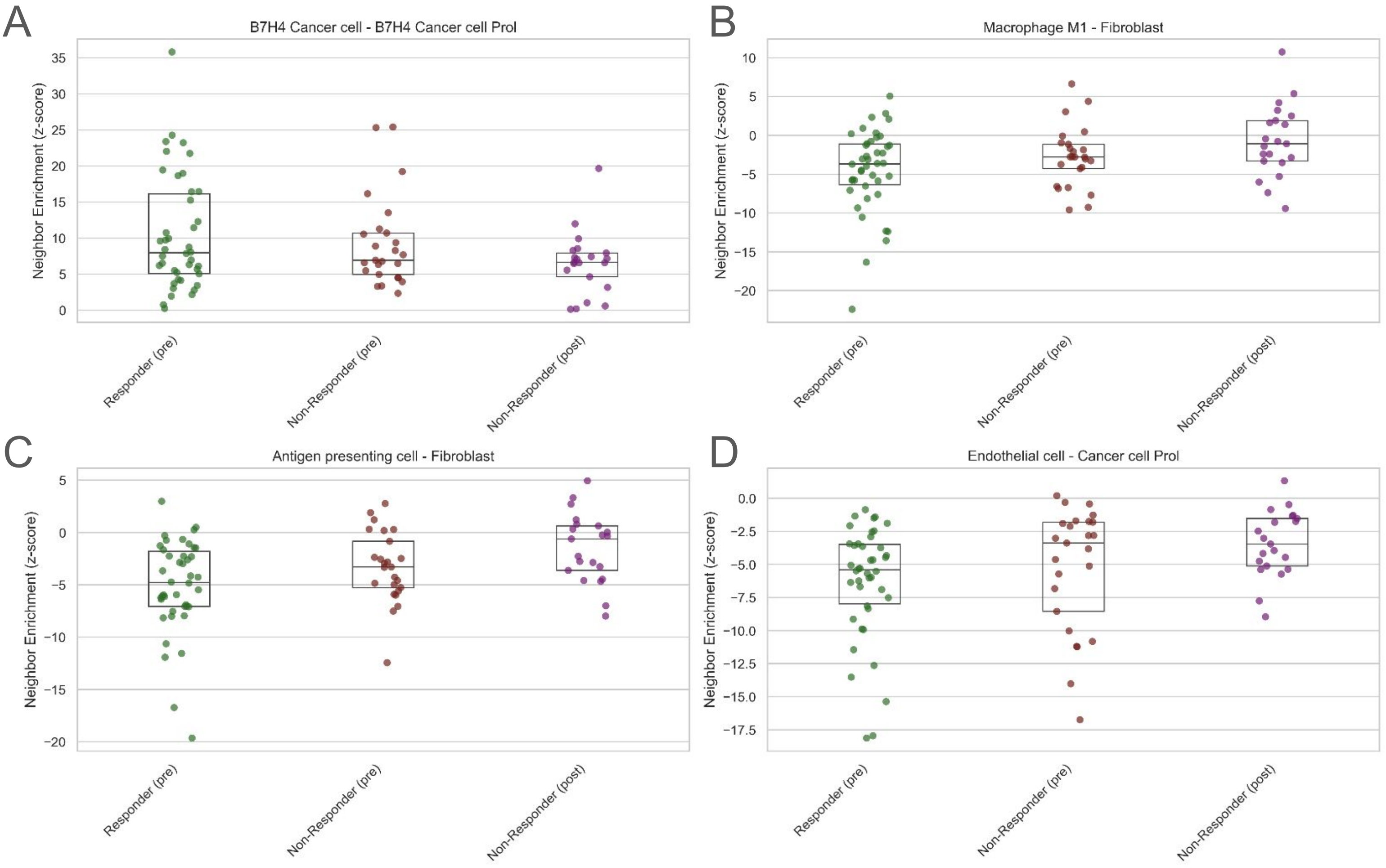


**Figure S5-2 (supplementary figure related to Figure 5)** Neighbourhood enrichment of selected pairs of cell types (see Results) in responders pre-treatment (green), non-responders pre-treatment (red) and non-responders post-treatment (purple). Each point is for a patient, showing the neighbor enrichment averaged across all ROIs of that patient.

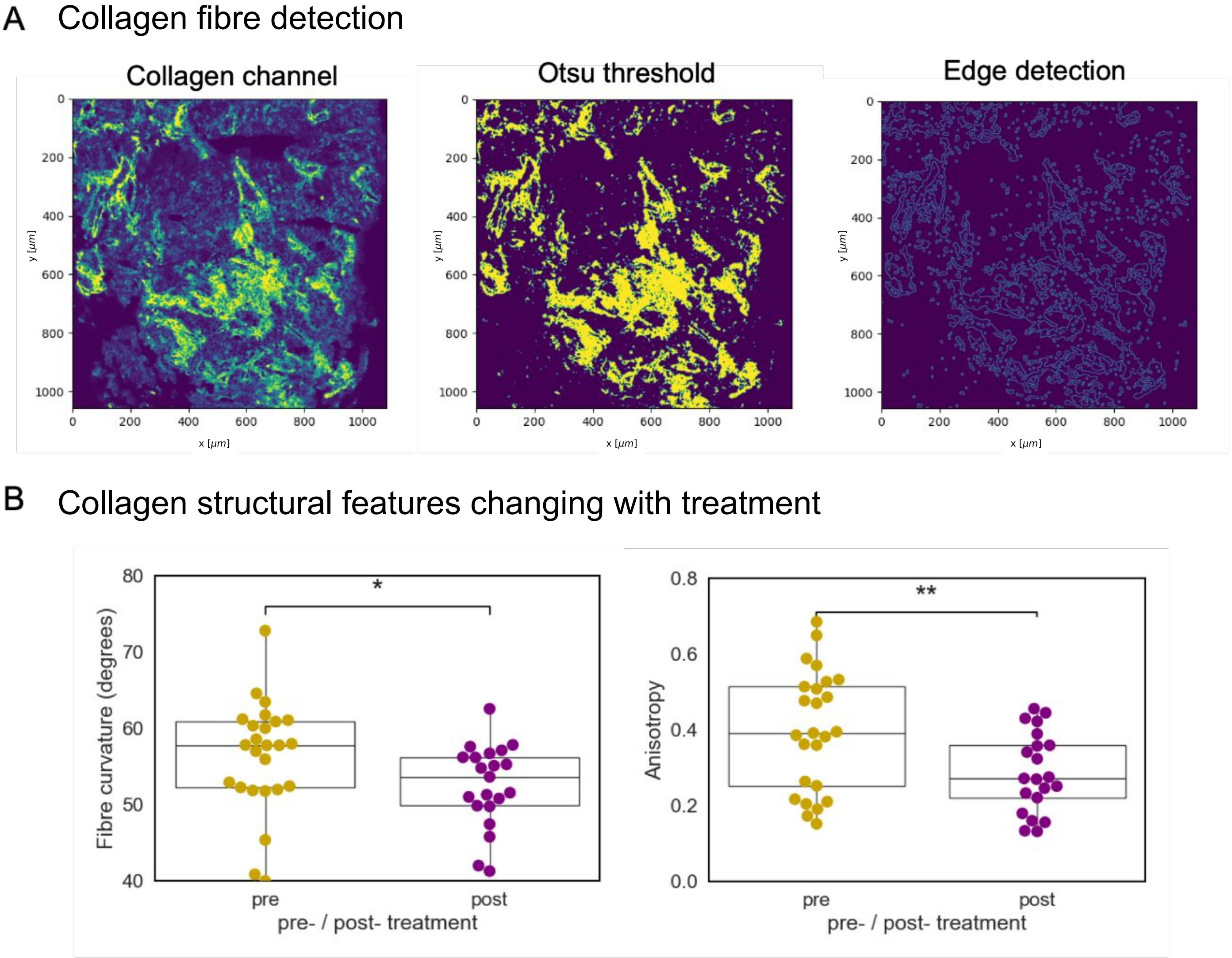


**Figure 6 - Collagen structure. A** Representative example of segmentation of a collagen image. (right) original pre-processed collagen image, (middle) thresholded image (right) application of the Canny filter. **B** Comparison of distributions of collagen features pre- and post-treatment for non-responder patients only. (left) Collagen fibre curvature (t-stat = -2.03, p-value = 0.049), (right) Anisotropy (t-stat = -2.82, p-value = 0.007). Boxplots display median, upper and lower quartiles.

A

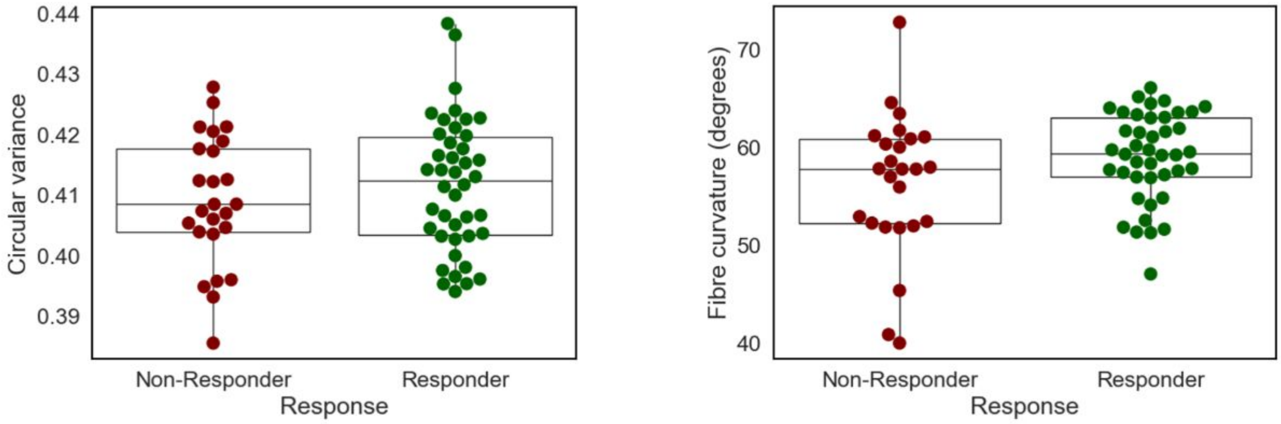


**Figure S6-1 (supplementary figure related to Figure 6) A** Selected collagen image features for responders (green) and non-responders (red). Circular variance of collagen fibres (left), t-stat = 0.938, p-value=0.353. Fibre curvature (right), t-stat=1.707, p-value=0.097.

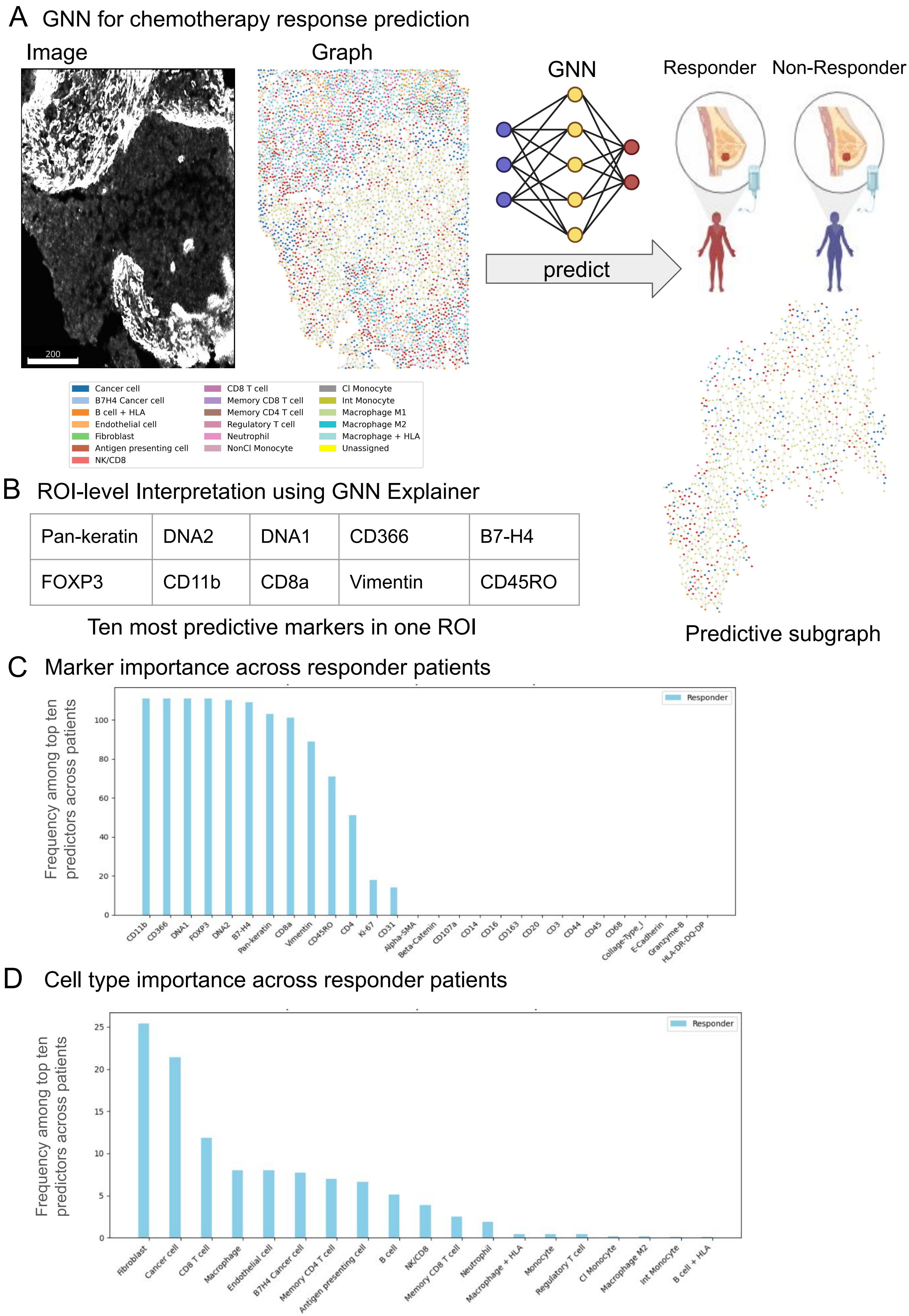
A GNN for chemotherapy response prediction
Image
Graph
GNN
Responder
Non-Responder
200
predict
Cancer cell
B7H4 Cancer cell
B cell + HLA
Endothelial cell
Fibroblast
Antigen presenting cell
NK/CD8
CD8 T cell
Memory CD8 T cell
Memory CD4 T cell
Regulatory T cell
Neutrophil
NonCl Monocyte
Cl Monocyte
Int Monocyte
Macrophage M1
Macrophage M2
Macrophage + HLA
Unassigned
B ROI-level Interpretation using GNN Explainer
Pan-keratin
DNA2
DNA1
CD366
B7-H4
FOXP3
CD11b
CD8a
Vimentin
CD45RO
Ten most predictive markers in one ROI
Predictive subgraph
C Marker importance across responder patients
Frequency among top ten predictors across patients
Responder
0
20
40
60
80
100
CD11b
CD366
DNA1
FOXP3
DNA2
B7-H4
Pan-keratin
CD8a
Vimentin
CD45RO
CD4
Ki-67
CD31
Alpha-SMA
Beta-Catenin
CD107a
CD14
CD16
CD163
CD20
CD3
CD44
CD45
CD68
Collage-Type_I
E-Cadherin
Granzyme-B
HLA-DR-DQ-DP
D Cell type importance across responder patients
Frequency among top ten predictors across patients
Responder
0
5
10
15
20
25
Fibroblast
Cancer cell
CD8 T cell
Macrophage
Endothelial cell
B7H4 Cancer cell
Memory CD4 T cell
Antigen presenting cell
B cell
NK/CD8
Memory CD8 T cell
Neutrophil
Macrophage + HLA
Monocyte
Regulatory T cell
Cl Monocyte
Macrophage M2
Int Monocyte
B cell + HLA

**Figure 7: Response prediction and interpretability. A** Graph neural network (GNN) model architecture integrating cell-cell contact graphs and protein abundance profiles to predict response to therapy. Each cell serves as a node with its 35-protein expression profile as features, connected by spatial proximity edges. Images on left show a representative IMC channel (scale bar is 200μm) and corresponding cell-cell contact graph, colored by cell type label (see legend). **B** Interpretability analysis for an example responder ROI using GNNExplainer yields a minimal predictive subgraph (right), and the ten protein markers contributing most to prediction performance (bottom). **C-D** Patient-level aggregation of protein marker importance scores (C) and cell type importance scores (D) associated with prediction of chemotherapy response, aggregated across all test samples from GNNExplainer analysis for each individual sample.
